# Comprehensive transcription factor perturbations recapitulate fibroblast transcriptional states

**DOI:** 10.1101/2024.07.31.606073

**Authors:** Kaden M. Southard, Rico C. Ardy, Anran Tang, Deirdre D. O’Sullivan, Eli Metzner, Karthik Guruvayurappan, Thomas M. Norman

## Abstract

Cell atlas projects have nominated recurrent transcriptional states as drivers of biological processes and disease, but their origins, regulation, and properties remain unclear. To enable complementary functional studies, we developed a scalable approach for recapitulating cell states *in vitro* using CRISPR activation (CRISPRa) Perturb-seq. Aided by a novel multiplexing method, we activated 1,836 transcription factors in two cell types. Measuring 21,958 perturbations showed that CRISPRa activated targets within physiological ranges, that epigenetic features predicted activatable genes, and that the protospacer seed region drove an off-target effect. Perturbations recapitulated *in vivo* fibroblast states, including universal and inflammatory states, and identified *KLF4* and *KLF5* as key regulators of the universal state. Inducing the universal state suppressed disease-associated states, highlighting its therapeutic potential. Our findings cement CRISPRa as a tool for perturbing differentiated cells and indicate that *in vivo* states can be elicited via perturbation, enabling studies of clinically relevant states *ex vivo*.

## Introduction

Ongoing cell atlas projects have revealed remarkable cellular diversity within the body, identifying over 500 distinct cell types^1–5^. Subsequent more targeted efforts within specific tissues have revealed that even greater diversity exists at the level of transcriptional state^6,7^, with some tissues like the brain comprising possibly thousands of transcriptionally distinct clusters^8^. Furthermore, many disease processes induce new transcriptional states, making them interesting candidates as drivers of pathology^9–11^. What is missing from these observational studies, however, is an understanding of how these transcriptional states came to be, how they are regulated, and what functional properties they may confer. Answering these questions *in vivo* is challenging because of the number of possible states, their rarity, the difficulty of targeting subpopulations with unclear origins, and the labor required to investigate each one individually.

Here we take a complementary approach by attempting to elicit these *in vivo* transcriptional states *in vitro* using genetic perturbations of transcription factors. This approach would enable simpler and more scalable experiments but requires tools for carefully manipulating gene expression in differentiated cells. While cDNA overexpression has been used to direct differentiation or force transdifferentiation^12–15^, it can lead to non-physiological expression levels or isoforms that may not optimally capture natural cell states. We hypothesized that CRISPR activation (CRISPRa)^16–18^, which mimics the regulatory events that naturally control gene expression, might be a more nuanced tool, given its past successes^19–25^ and the notable enrichment of transcriptional regulators in the hits of prior screens^16,26^. However, two challenges arise in this endeavor: the need to be sensitive to the many possible outcomes of transcription factor perturbations, and persistent issues with CRISPRa guide efficacy^25,27,28^. Perturb-seq, like other single-cell CRISPR screens^28–38^, offers a solution by linking CRISPR-mediated genetic perturbations to their phenotypic effects via single-cell RNA sequencing. A single experiment can therefore enable simultaneous investigation of the drivers of multiple transcriptional states and the direct characterization of on-target activity.

Our initial target is the fibroblast, a phenotypically plastic cell type that is ubiquitous in the body as the workhorse of connective tissue. In response to injury, fibroblasts become activated, leading to marked changes in morphology and behavior as they help orchestrate wound healing. Pathological activation results in fibrosis, a common cause of death with few treatments^39^. They are also mediators of inflammation in diseases like rheumatoid arthritis^40–42^, and major constituents of the tumor microenvironment^43^. Perhaps due to their natural plasticity, cell atlas studies revealed substantial fibroblast transcriptional heterogeneity, with some states tied to phenotypic differences^41,42,44–54^. A large-scale study^53^ across 17 tissues identified unique fibroblast subpopulations in mice and humans, including an *Lrrc15*^+^ state in cancer-associated fibroblasts and a *Dpt*^+^ *Pi16*^+^ “universal” fibroblast state present in most tissues, hypothesized to be immunomodulatory and to serve as a progenitor state for other subtypes^50^. Another study^54^ identified 17 clusters of human fibroblasts across inflammatory diseases, and found likely concordant states: a *SPARC*^+^ *COL3A1*^+^ inflammatory state that was expanded in all diseases, and a *CD34*^+^ *MFAP5*^+^ state similar to the universal state. These findings suggest there is potential therapeutic value in promoting possibly beneficial states, such as the universal state, or suppressing disease-associated states, such as the inflammatory state, by identifying their respective regulators.

Here we perform large-scale CRISPRa Perturb-seq experiments to test our ability to elicit these and other fibroblast transcriptional states. We exploit a simple and widely applicable modification to increase the throughput of Perturb-seq based on droplet overloading, enabling screens targeting 1836 candidate transcription factors with 10,979 guide RNAs in primary human fibroblasts and retinal pigment epithelial cells. These experiments revealed key principles governing CRISPRa activity and specificity, most notably that it appears to activate transcription factors within a physiological expression range, establishing it as a nuanced tool for manipulating the transcriptome of differentiated cell types. Furthermore, our experiments identify key regulators driving the emergence of distinct fibroblast transcriptional states, including the universal and inflammatory phenotypes, and reveal cross-regulation between them that may reveal new therapeutic targets. Taken together, these results highlight the power of CRISPRa Perturb-seq to understand and recapitulate *in vivo* transcriptional states.

### Droplet overloading for high-throughput Perturb-seq

To enable unbiased studies, we set out to replicate a cDNA library targeting 1,836 candidate transcription factors^13^ using CRISPRa as the overexpression method. We constructed a VPH-dCas9 CRISPRa effector (**Figure S1A,** Methods)^17,21^ that showed robust and uniform activation of cell surface markers (**Figure S1B**). However, guide selection posed a challenge, as predicted CRISPRa guides from existing libraries were often inactive: e.g., only 3/8 guides targeting *CD45* showed activity (**Figure S1C**). To ensure redundancy, we therefore chose 6 guides per target from existing libraries^26,55–57^, with rigorous off-target filtering (Methods). Our final library comprised 10,979 guides, including 78 overrepresented non-targeting negative control guides (**Figure S1D**, **Supplemental Table 1**).

Performing experiments at this scale with existing techniques was prohibitively expensive. This challenge has motivated the development of multiplexing strategies that trade off single-cell resolution for the ability to measure the average effects of more perturbations. These strategies include introducing multiple guides per cell by superinfection^58–60^ or “overloading”^59,61^ scRNA-seq assays so that more droplets contain cells, including more multiplets (2+ cells per droplet) (**Figure 1A**). In both cases, the average effects of perturbations can be reconstructed by solving a regression problem to demultiplex the superimposed effects (**Figure 1B**).

**Figure 1.**
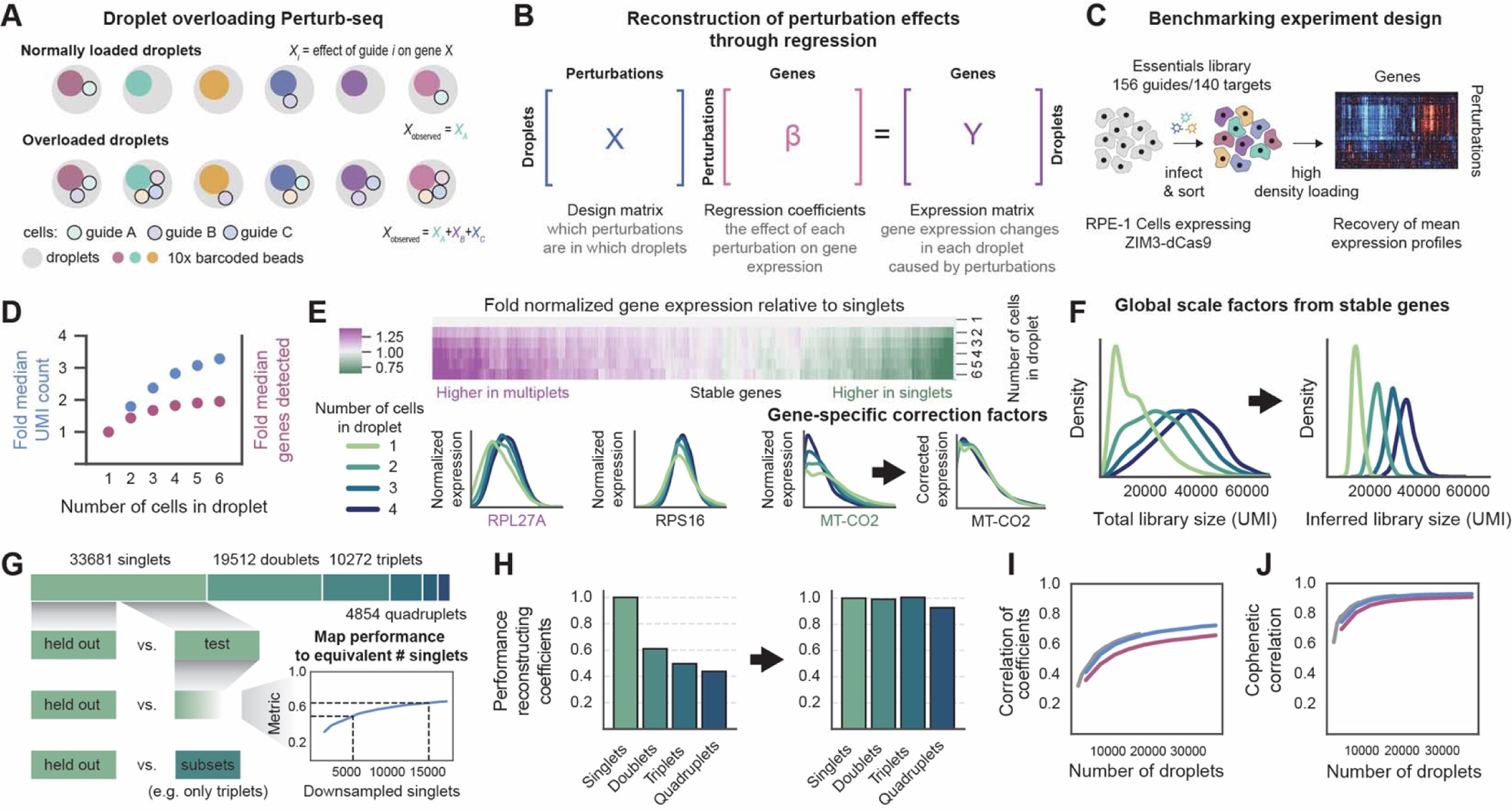
Droplet overloading for high-throughput Perturb-seq. A. Schematic of the droplet overloading approach: increased cell loading leads to more multiplets (2+ cells), enabling measurements of more perturbations by trading off single-cell resolution. B. Regression can be used to demultiplex superimposed perturbation effects in overloaded droplets. C. Design of benchmarking experiment. D. Single-cell RNA sequencing library size and complexity as droplet occupancy increases. E. Variability in transcript capture efficiency as a function of droplet occupancy. Some transcripts like *RPL27A* or *MT-CO2* are captured more efficiently in multiplets or singlets, respectively, while stable genes are captured similarly across conditions. Gene-level corrections (bottom right) are fit to normalize for these differences. F. Fitting global scale factors to stably captured genes results in estimates of true library size that are better resolved by droplet occupancy. G. Overview of the benchmarking approach used to assess reconstruction accuracy. Results from disjoint subsets of the data are compared. Baseline performance is determined from comparing equal numbers of singlets. By downsampling one set, performance of other subsets can be quantified in terms of the equivalent number of singlets required to achieve the same reconstruction accuracy. H. Reconstruction accuracy (relative to equivalent number of singlets) using only singlets, doublets, triplets, or quadruplets, with and without normalization. I. Correlation between expression profiles measured using up to 16,800 singlets (grey) or reconstructed using varying numbers of droplets (each containing up to 6 cells). Blue and magenta denote performance with and without normalization, respectively. J. Same analysis as (I) but compared using the correlation of correlation matrices, which capture functional relationships among perturbations.

We focused on the overloading approach because it separates perturbations across distinct cells, preventing unmodeled interactions and ensuring that strong perturbations do not overwhelm weak ones. This approach is particularly beneficial for studying transcription factor perturbations, which are likely to interact if introduced at high multiplicity. To develop our approach, we constructed a library of 155 single guide RNA (guides) targeting 140 essential genes (plus 16 non-targeting controls) that clustered in a previous screen (**Figure 1C, Figure S2A, Supplemental Table 1**)^62^. We transduced this library into RPE-1 cells stably expressing a ZIM3-dCas9 CRISPRi effector construct at low multiplicity of infection (MOI), sorted sgRNA-containing cells by fluorescence assisted cell sorting (FACS), and loaded ∼125,000 cells per lane onto two lanes of the 10x Genomics Chromium to achieve overloading (approximately eight times the typical cell loading density). The resulting scRNA-seq libraries contained ∼36,000 loaded droplets per lane, split evenly between singlets and multiplets.

We observed a saturation effect as droplet occupancy increased (**Figure 1D**) that is not accounted for in existing methods^59^. For example, scRNA-seq libraries made from doublet-containing droplets had only ∼1.8X as many total RNA molecules (measured in unique molecular identifiers, UMIs) as from singlet-containing droplets. Library complexity (median detected genes) nevertheless continued to increase, suggesting library construction was not inhibited by overloading. Interestingly, we found that saturation was transcript-dependent. Estimates of average transcript capture efficiency for droplets of different occupancy levels revealed highly variable patterns, with some transcripts like *RPL27A* being captured more efficiently in multiplets, while others such as *MT-CO2* (and other mitochondrial transcripts) exhibited declining capture efficiency as droplet occupancy increased (**Figure 1E**).

To compensate for these saturation effects, we developed a new computational normalization (Methods). First, we introduced gene-level scaling factors that equalize capture rates across occupancies and identified “stable” genes whose capture rate was largely invariant (**Figure 1E**, center). For each droplet, we fit a global scale factor to these genes to infer the library size if all genes were stably captured; these estimates were better resolved by occupancy than the observed library sizes (**Figure 1F**). To normalize, we applied the gene-level corrections, scaled by the inferred library size, and centered and scaled each gene by its mean and standard deviation in control cells (Methods). With this saturation-aware normalization, average phenotypes could be reconstructed from all droplets using linear regression (**Figure 1B**).

We benchmarked the reconstructions by comparing replicability between different subsets of the data (**Figure 1G**). As a baseline, we split the singlets into two equal sets, independently computed average effects, and downsampled one set to create a calibration curve mapping reconstruction accuracy to the number of singlets that would yield equivalent performance (Methods). We first compared (using Pearson correlation, **Figure S3A**) the reconstructed average phenotypes using singlets versus doublets, triplets, or quadruplets. Without saturation-aware normalization, performance dropped strongly for higher occupancies (**Figure 1H** left). With saturation-aware normalization, we observed approximately one-to-one performance by droplet number: e.g., 8,400 doublets yielded equivalent results to 8,400 singlets (**Figure 1H** right). Similar behavior was seen using varying numbers of doublets (**Figure S3B**) or all droplets containing up to 6 cells (**Figure 1I, S3C**). This normalization significantly improved efficiency, as without it, two to three times as many multiplets were needed to match the performance of conventional Perturb-seq (**Figure 1I**). Average phenotypes inferred from saturation-aware normalized droplets were visually indistinguishable from those using only singlets (**Figure S3D** top), whereas structural differences were apparent without saturation-aware normalization (**Figure S3D** bottom). As an alternative metric, we assessed how well the correlations between perturbations, which can drive clustering and functional insights, were recovered. Performance was strong (**Figure 1J**) and improved by saturation-aware normalization (**Figure S3E**).

Finally, significant regression coefficients mark differentially expressed genes in this procedure. We observed improved stability in identifying differentially expressed genes (**Figure S3F-H**), and a consistent increase in the number detected per perturbation when using all droplets compared to all singlets (**Figure S3I**). Our approach therefore provided a simple and scalable means to conduct large Perturb-seq experiments by correcting the technical biases introduced by overloading.

### Assessing strength and determinants of CRISPRa on-target activity using Perturb-seq

We next applied our droplet overloading Perturb-seq approach to activate all transcription factors in two untransformed cell lines: Hs27 primary foreskin fibroblasts and RPE-1 retinal pigment epithelial cells. In total we measured about 808,000 and 1.76 million cells in Hs27 and RPE-1, respectively (**Figure 2A**, **Figure S2B-C**, Methods), across ten 10x HT lanes each and with even representation across perturbations (**Figure S2B,C**).

**Figure 2.**
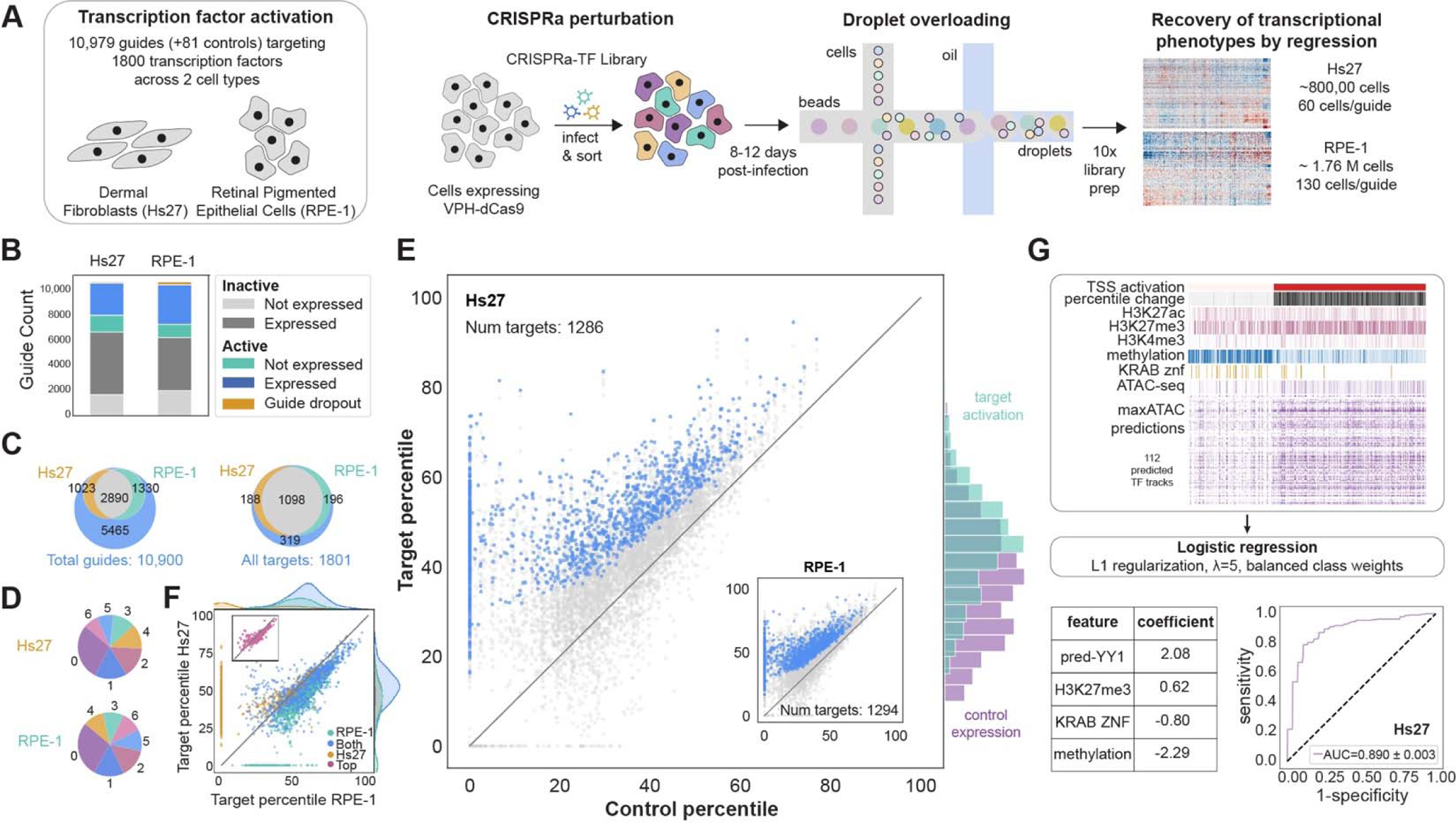
Efficacy of CRISPRa against transcription factors measured using Perturb-seq. A. Overview of large-scale transcription factor CRISPRa Perturb-seq experiments. B. Observed on-target activity for all guides according to whether target gene is expressed. C. Overlap of guide activity across two cell types. Left shows number of guides that are inactive (blue), active only in Hs27 (yellow), active only in RPE-1 (green), and active in both (grey). Right shows the number of targets that are inactivable (blue), activable in Hs27 (yellow), activable only in RPE-1 (green), and activable in both (grey). D. Number of active guides per target gene. E. Distribution of induced expression changes by CRISPRa for Hs27 and RPE-1 (inset). Expression of a target gene is quantified by the percentile rank relative to all genes in control cells. Best performing guides for each target are shown in blue. Right histograms compare induced expression of unexpressed target genes to distribution of expressed transcription factors in control Hs27 cells F. Comparison of target activation across cell types. Guides showing activation across cell types are shown in blue, while yellow and green indicate those which showed specific activation in Hs27 and RPE-1 respectively. Inset (red) compares only best performing guides for each target gene. G. Epigenetic determinants of susceptibility to CRISPRa. Top: Schematic of the model building approach used to identify epigenetic features predictive of susceptibility to CRISPRa. Bottom left: Selected features and coefficients of model fit in Hs27 cells. Bottom right: Performance of model.

These large datasets enabled direct assessment of on-target efficacy for 21,958 CRISPRa perturbations. The low expression of some transcription factors prevented us fitting the gene-level correction factors previously described, so we developed a simplified regression approach to quantify activation (Methods). Despite >50% of guides appearing inactive (**Figure 2B**, **Figure S4A,B**), 1,482 transcription factors could be activated in at least one cell type, including targets that were unexpressed according to baseline bulk RNA sequencing (**Figure 2C**, **Figure S4C**, **Supplemental Table 4**). Most activatable targets had multiple active guides (**Figure 2D**), suggesting a low false negative rate, and most (80%) expressed genes had at least one active guide.

cDNA overexpression approaches typically express all genes with a common, often strong, exogenous promoter. In contrast, little is known about the range of expression levels induced by CRISPRa. To enable comparisons in a common scale, we quantified a gene’s expression in terms of its percentile rank relative to all genes in control cells (Methods); e.g., CRISPRa might activate a target from the 20^th^ expression percentile to the 50^th^ percentile. The top guides for each target consistently increased expression above control levels (**Figure 2E** and inset, blue dots), with a minority of guides causing expression to decline. On-target activity was also well-correlated across cell types (**Figure 2F**), particularly for top guides (**Figure 2F**, inset).

Critically, CRISPRa activated unexpressed targets to a similar expression range seen in endogenously expressed transcription factors in control cells (**Figure 2E**, right), suggesting its activity is constrained by the endogenous gene regulatory environment. Accordingly, there were diminishing returns when trying to further activate already expressed genes (**Figure S4D**). CRISPRa-mediated activation of a target also yielded a similar expression level to the maximal induction seen as an indirect result of activating another transcription factor (**Figure S4E,F**).

Constraints were also observed in terms of which genes could be activated, with cell-type-specific activation for 188 (Hs27) and 196 (RPE-1) targets (**Figure 2C** right, **Figure S4C**) and 319 genes resistant to activation entirely. Our dataset therefore provided a unique opportunity to explore the determinants of susceptibility to CRISPRa for many endogenous genes that were unexpressed in control cells (**Figure S4G**). Previous work has shown epigenetic context plays a role^63^, so we performed ATAC-seq and CUT&RUN assays targeting histone modifications (H3K27ac, H3K27me3, and H3K4me3) on both Hs27 fibroblast and RPE-1 cells. We focused our initial analysis on Hs27 cells, to incorporate publicly available skin fibroblast DNA methylation data^64^. We also used maxATAC^65^ to predict binding activity for 112 Hs27-expressed transcription factors from our ATAC-seq data. Finally, we observed that KRAB zinc finger proteins were resistant to activation (**Figure S4H**), so we created a feature based on membership in KRABopedia^66^. In total, we collected 118 features, which we summarized for each target gene within a window centered at the transcription start site (TSS, Methods).

To identify features that were predictive of susceptibility to CRISPRa, we performed penalized logistic regression (**Figure 2G, top,** Methods), which selected four features: H3K27me3, KRAB zinc finger identity, DNA methylation, and maxATAC-predicted YY1 binding (pred-YY1) (**Figure 2G, bottom left**). A model trained using these features achieved a 5-fold cross-validated AUC of 0.89 (**Figure 2G, bottom right**), with the features exhibiting clear differences at target promoters according to susceptibility to activation (**Figure S4I**). The model could also distinguish between cases where targeting one of two TSSes for a gene resulted in activation while targeting the other did not (**Figure S4J,K**). pred-YY1 and DNA methylation showed the strongest positive and negative effects, respectively, and models trained only with individual features confirmed these had the strongest predictive power (**Figure S4L**). Notably, pred-YY1 outperformed the ATAC-seq signal. The second strongest positive signal, aside from ATAC, was H3K27me3, consistent with Enrichr^67,68^ enrichment analysis of activatable genes using the “ENCODE_Histone_Modifications_2015” gene set (**Figure S4M**).

We then tested the generality of our model by exploring whether the model trained on Hs27 cells could predict CRISPRa activity in RPE-1 cells. Because DNA methylation data was unavailable, we trained a reduced model using three features: pred-YY1, H3K27me3, and KRAB zinc finger identity. When evaluated on RPE-1 data, this reduced model achieved an AUC of 0.883 (**Figure S4N**). Consistent with Hs27 cells, activated promoters in RPE-1 showed high pred-YY1 (**Figure S4I, bottom**). The reduced model also captured cell-type-specific differences in CRISPRa susceptibility, as evidenced by higher predicted activation probabilities for 33 genes uniquely activated in Hs27 cells compared to RPE-1 cells (**Figure S4O**).While these findings leave open the role of YY1 in mediating gene activation by CRISPRa, they reveal that the susceptibility of unexpressed genes to activation by CRISPRa is a largely predictable product of epigenetic context at target promoters, and that activation is constrained to a physiological expression range.

### Seed-dependent off-target effects of CRISPR epigenetic editors

The 21,958 transcriptional phenotypes we gathered represent one of the largest high-content CRISPR perturbation datasets gathered to date. Critically, each cell received only a single guide RNA and we targeted each gene with six independent guides across two cell types. This scale and the degree of internal replication within our experiment allowed us to investigate structural patterns in CRISPRa’s activity in altering the transcriptome that were not previously possible. We applied our regression approach to reconstruct average transcriptional phenotypes for all perturbations and uncovered two key features. First, and as expected, guides inducing strong target activation did not necessarily induce strong transcriptional phenotypes (**Figure S5A**), likely reflecting context-dependent activity or regulation of transcription factors. Second, despite rigorous filtering during guide library design (Methods), clustering revealed an off-target effect: 9.7% of guides in Hs27 cells and 26.4% in RPE-1 cells clustered into large groups of guides sharing the same seed region (∼5 PAM-proximal nucleotides, **Figure 3A, right**).

**Figure 3.**
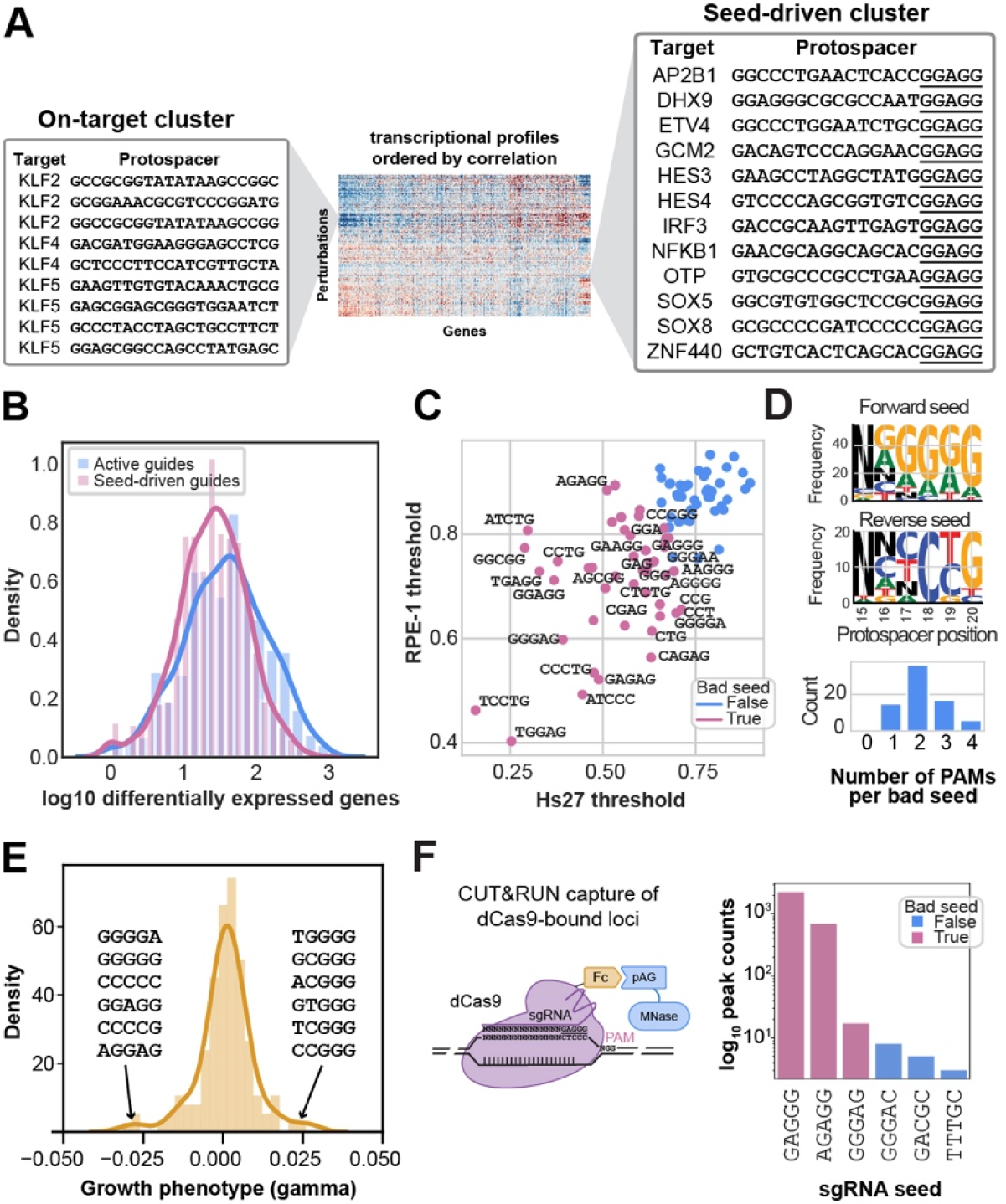
Off-target effects driven by the seed region of the protospacer. A. Schematic depicting perturbation profile clustering driven by seed region of the protospacer in RPE-1 cells. Many guides formed clusters with other guides with the same seed region of the protospacer (underlined). B. Comparison of the number of differentially expressed genes induced by on-target activity vs. seed-based off-target activity in Hs27. C. Common seeds driving clustering across Hs27 and RPE-1 experiments. Lower thresholds correspond to “bad” seeds that drive clustering. D. Enrichment of PAM sequences in seeds driving clustering. Top: We categorized bad seeds as either G-rich or C-rich and then computed sequence logos. Bottom: Number of PAM sequences within bad seeds. E. Median fitness effects in non-targeting control sgRNAs grouped by seed in a genome-wide CRISPRa fitness screen, Gilbert *et al*., 201416. F. CUT&RUN capture of Fc-dCas9-bound loci to profile seed-driven guide binding across the genome.

Seed-dependent differences in Cas9 binding in mammalian cells^69^ and seed-dependent off-target effects of CRISPRi in bacteria^70^ have been reported. Here we found that CRISPRa seed-driven effects could be as strong as on-target effects in terms of overall impact (**Figure 3B**). We identified a common set of seeds that drove clustering across the two cell types (**Figure 3C**), which were notably enriched for PAM sequences (NGG/NAG) or their reverse complements (CCN/CTN) (**Figure S3D**). The presence of a “bad” seed did not hinder on-target activation (**Figure S5B**), suggesting that guides direct a combination of specific on-target effects driven by the full protospacer sequence and off-target effects driven by the seed. Accordingly, we could regress out apparently similar seed-driven effects in previous genome-wide CRISPRa fitness screens^16,26,71^ (**Figure S5C-F,** Methods), including a case in which the median fitness phenotypes of non-targeting negative control guide RNAs, grouped by seed, identified similar PAM-rich sequences (**Figure 3E**)^16^. In contrast, a CRISPRi fitness screen did not show obvious enrichment for PAM-containing seed sequences (**Figure S5G**), although we could identify seed-driven clusters in a previous large-scale CRISPRi Perturb-seq experiment (**Figure S5H**). Finally, a CRISPRoff fitness screen^72^ showed signs of seed-driven effects (**Figure S5J**).

To directly assess off-target binding, we developed a dCas9 CUT&RUN protocol (Methods). Profiling of six guides revealed that bad seeds led to more binding sites within the genome **(Figure 3F**). Seed-driven effects therefore likely arise from actions of Cas9 epigenetic effectors at these alternate binding sites. We incorporated both our on-target effect activity estimates and seed filtering to produce a library of validated active sgRNAs for 1319 transcription factors (**Supplemental Table 4**).

### Expression programs and regulatory interactions driven by transcription factors in fibroblasts

We next examined the global architecture of transcription factor-induced phenotypes. Activating individual transcription factors had more subtle effects than, for example, knockdown of essential genes^62^. To account for this difference and to ensure resilience against off-target effects, we developed a clustering-based test to identify transcription factors that robustly altered the transcriptome (**Figure 4A**, top right). This method leverages the principle that when the dataset is clustered at a sufficiently fine resolution, guides targeting the same gene are unlikely to co-cluster by chance; instead, clustering likely reflects a common transcriptional effect from target gene activation. This test assesses both reproducibility (multiple guides must cluster) and distinctiveness (clusters must emerge) of induced transcriptional phenotypes. Clustering by profile correlations, which is robust to guide efficacy differences^28,62^, and applying this criterion identified 212 and 178 transcription factors with strong effects in Hs27 and RPE-1 cells, respectively, plus 102 and 82 additional genes that had single guides that clustered strongly with these gold standards that we included in our analyses (Methods). In total, 1077 guides targeting 417 genes induced a strong transcriptional phenotype in at least one cell type (**Figure S6A**).

**Figure 4.**
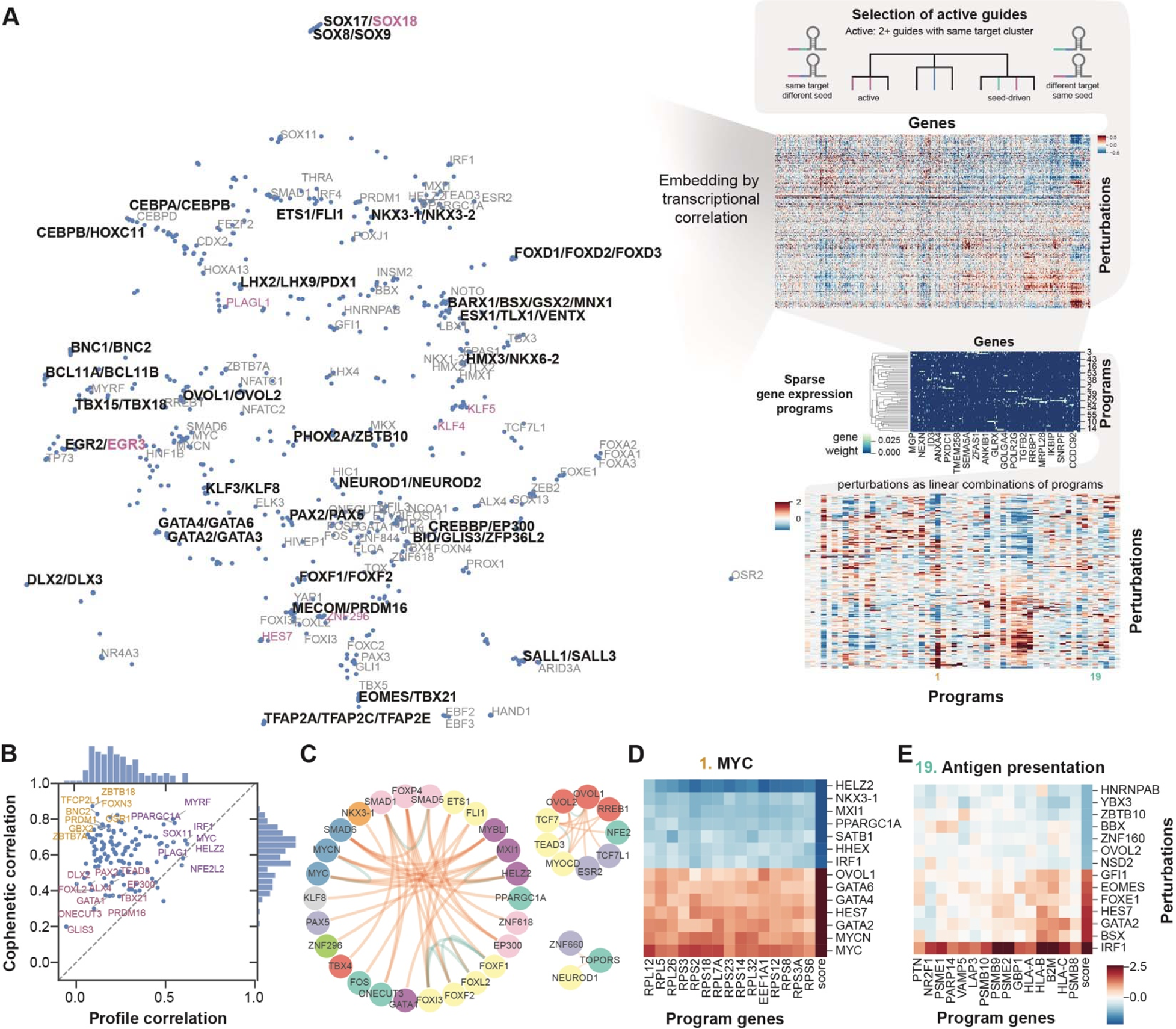
Expression programs and regulatory interactions driven by transcription factors in fibroblasts. A. Overview of the approach used to identify interpretable transcription factor-driven gene expression programs. Transcriptional phenotypes are reconstructed for all perturbations and a clustering-based test is applied to identify robust hits attested to by multiple guides. Sparse PCA is then used to extract positively correlated gene programs from these phenotypes. Embedding shows hit transcription factors for Hs27 fibroblast cells. Some hits cluster tightly together, indicated by slashes in labels. Magenta marks transcription factors highlighted in later analyses. B. Comparison of transcription factor effects between Hs27 and RPE-1 cells using two metrics: direct correlation of induced expression changes (*x*-axis) vs. correlation of perturbation-perturbation correlation matrices (*y*-axis). C. Anticorrelated perturbation pairs suggestive of regulatory interactions. Arcs in the circos plots annotate strong negative (red) or positive associations (teal) among the selected transcription factors. D. The top activators (red) and repressors (blue) the of the broadly modified *MYC* gene program. E. The top activators (red) and repressors (blue) the of the highly IRF1 perturbation specific antigen presentation program.

We visualized relationships among hit guides by embedding their transcriptional phenotypes (Hs27: **Figure 4A**, RPE-1: **Figure S7**, Methods). Guides clustered primarily by target and subsequently by homology, as evidenced by clustering of genes with similar names. For instance, guides targeting *BCL11A*, *BCL11B*, and *ZNF827*, which share an amino-terminal motif that mediates NuRD complex interactions^73^ clustered in both datasets, as did the *CREBBP*/*EP300* complex. However, most clusters were cell-type-specific: for example, *CRX* and *OTX2* are critical regulators of retinal development^74^, and clustered only in RPE-1 cells (**Figure S7**). To examine how transcription factors’ effects varied across cellular contexts, we compared the effects of 121 perturbations active in both cell types using two approaches: by directly correlating the expression profiles, and by correlating the correlation matrices of the perturbations, which assesses similarities in how perturbations relate to each other (Methods). While similar expression changes (**Figure 4B**, top right) were rare and typically associated with specialized inducible regulons, the relationships between transcription factors were better preserved (**Figure 4B**, top left), suggesting regulatory interactions are conserved despite differences in how effects manifest. To identify strong interactions, we examined anticorrelated perturbation pairs (**Figure 4C**, Methods). This analysis revealed known interactions, such as *MYC* repression by *HELZ2*^75^ and *MXI1* and *SMAD5* inhibition by *SMAD6*, as well as many candidate interactions.

To further understand how these transcription factors orchestrated gene expression, we sought to identify modules of functionally related genes that they regulated. We developed a sparse PCA approach (**Figure 4A**, right middle, Methods) to identify reproducible and interpretable gene expression programs (Methods). Applied to Hs27 cells, this procedure organized 2,389 genes into 57 programs (**Figure 4A**, bottom right, **Figure S8, Figure S9A**, Supplemental Table 2). Though the programs were sparse, transcription factors usually modified multiple programs simultaneously. Nevertheless, perturbations often showed a distinct preference for specific programs. For example, one program (**Figure 4D**) was dominated by translation-associated genes, was strongly induced by *MYC* and *MYCN*, and was strongly suppressed by *MYC* repressors like *MXI1* and less characterized modulators like *HELZ2*^75^ and *NKX3-1*^76^, suggesting it reflects *MYC* activity. Other broadly activated programs corresponded to transcriptional responses like the cell cycle and mTORC1-mediated activation of *ATF4* (**Figure S9B,C**), while some were highly specific, such as an *IRF1*-driven antigen presentation program (**Figure 4E**) and an oxidative stress response driven largely by *NFE2L2* (**Figure S9D**).

### Transcription factor perturbations driving markers of *in vivo* fibroblast states

As our transcription factor perturbations were physiological and exhaustive, we reasoned that some gene programs might explain fibroblast transcriptional heterogeneity observed *in vivo*. We performed an enrichment analysis (Methods) to test whether gene programs overlapped with differentially expressed genes associated with states in a human cross-tissue stromal atlas^54^. We found significant overlaps for several gene programs (**Figure S10A**).

The first program corresponded with the *MYH11^+^ in vivo* state, included canonical smooth muscle markers such as *ACTA2*, *CNN1*, and *TAGLN*^77^, and captured transcription factors promoting a myofibroblast phenotype^49^ (**Figure 5A**). Top activators included the smooth muscle regulator *MYOCD*^78,79^, the pro-fibrotic Hedgehog signaling effector *GLI1*^80,81^, and regulators of epithelial-to-mesenchymal transition (EMT) such as *SNAI1* and *FOXC2*^82^. Conversely, *FLI1* and *KLF5*, whose reduced expression is associated with systemic sclerosis^83^, were negative regulators. Many novel inducers emerged, such as the Notch effector *HES7* (**Figure 5A**, magenta), known for its role in somitogenesis^84^, and the uncharacterized gene *ZNF296* (**Supplemental Table 2**). Guides targeting these genes upregulated *ACTA2* and induced myofibroblast-like morphological changes (**Figure 5B**, **Figure S10B**), with the *HES7* results consistent with the role of Notch signaling in controlling fibroblast state^40,41^. Interestingly, *EOMES*, a transcription factor that regulates EMT in a sharply timed period of early development^85^, was also a strong inducer. Our perturbative approach can therefore be used to study phenotypes that are rare relative to typical expression patterns.

**Figure 5.**
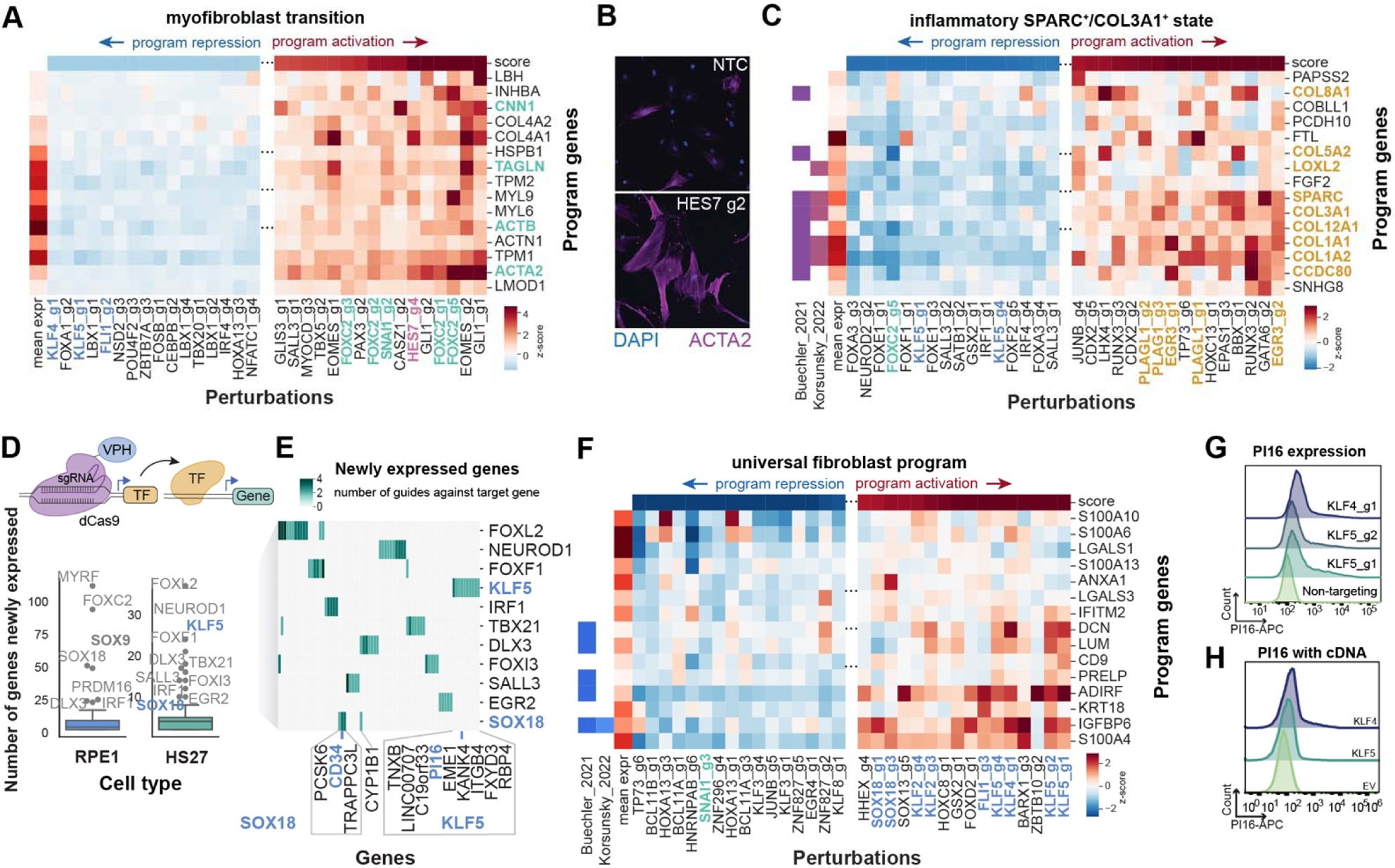
Transcription factor perturbations driving markers of *in vivo* fibroblast states. A. Identification of a myofibroblast gene expression program and its regulators. Top genes defining the program form the rows of the heatmap, and top perturbations repressing (left) or activating (right) the program form the columns. Green denotes canonical marker genes or perturbations while magenta marks transcription factor hits highlighted in further analysis. B. Morphological changes and ACTA2 induction by *HES7*. C. Identification of an inflammatory fibroblast gene expression program and its regulators. *In vivo* inflammatory state marker genes, including *COL1A1, COL1A2, COL3A1, COL5A2, SPARC*, are colored yellow with highlighted perturbations sharing the same color. Genes exhibiting overlap with inflammatory state genes in references 53 and 54 respectively are indicated (left). D. To score the ability of perturbations to induce new expression states, we counted how many genes they activated that were unexpressed in control cells. Labeled points in the box plots show outlier perturbations for each cell type. E. Examples of genes newly activated by top Hs27 hits from this analysis. Plot is colored according to how many independent guides led to activation. Marker genes *PI16* and *CD34*, associated with the universal fibroblasts, are highlighted. F. Perturbations that induced markers of universal fibroblasts (see E) also activated a sparse gene expression program. Genes exhibiting overlap with universal state genes in references 53 and 54 respectively are indicated (left). G. Flow cytometry validation of PI16 induction by predicted regulators using CRISPRa. H. Flow cytometry validation of PI16 induction by predicted regulators using cDNA overexpression.

A second program corresponded with the *SPARC^+^COL3A1^+^*inflammatory fibroblast population (**Figure 5C, Figure S10A**)^54^. Although we did not observe *LRRC15* expression in our experiment, the identified program included the genes *COL1A1*, *COL1A2*, *COL3A1*, *COL5A2*, and *SPARC* that are upregulated in the cross-tissue atlas *Lrrc15*^+^ mouse fibroblasts^53^, *LRRC15*+ cancer associated fibroblasts (CAF)^86^, and within a *Lrrc15*^low^ CAF cluster in a model of pancreatic ductal adenocarcinoma^52^. Numerous perturbations altered the expression of this gene program in both directions. Strong inducers included *PLAGL1*, *PLAG1*, *GATA6*, and *EGR3*, with the latter two being previously reported as possible mediators of fibrogenic responses^87,88^. Conversely, strong suppressors included *NEUROD2*, *FOXE1*, *FOXA3*, and *KLF5*. According to the Human Protein Atlas^2^, the main suppressors of the program, aside from *KLF5*, have much stronger expression in non-mesenchymal cell types, suggesting their expression may alter core fibroblast function. In contrast, the main inducers of the program are expressed ubiquitously.

Notably, *KLF5* suppressed each of these programs. It also scored highly in a second analysis we conducted aimed at identifying transcription factors that elicited entirely new transcriptional states. We measured this by counting the number of genes each perturbation activated across two or more independent guides that were previously unexpressed in control cells (**Figure 5D**, Methods). Applied to RPE-1 cells, this approach for example identified the established pioneer factor *SOX9*^89^ (**Figure 5D, left**), which activated glial markers including *S100B* and *KDR*^2,90^ (**Figure S10C**). Remarkably, in Hs27 cells, perturbation of *KLF5* and *SOX18* induced expression of the universal fibroblast markers *PI16* and *CD34*, respectively^53,54^ (**Figure 5E)**. We expanded our search (**Figure S10D**) by assembling a curated list of fibroblast transcriptional state markers from the literature (**Supplemental Table 3**) and including all perturbations showing clear on-target activity (Methods), revealing induction of universal fibroblast markers by homologs of our initial hits, such as *CD34* by *SOX17* perturbation and *TNXB*^53^ by *KLF4* and *KLF5*. Finally, *KLF4*, *KLF5*, and *SOX18* all scored as inducers of a gene program associated with the *in vivo CD34^+^MFAP5^+^* state (**Figure 5F, Figure S10A**), which we termed the universal fibroblast program.

To validate these phenotypes, we constructed Hs27 cell lines individually expressing all guides that showed evidence of *PI16* induction (including those previously filtered due to low on-target activity)— *KLF4*, *KLF5*, and *ASCL5*. Flow cytometry analysis confirmed that all selected guides induced PI16 production in an apparently bimodal pattern (**Figure 5G**, **Figure S10E**), with similar phenotypes obtained via cDNA overexpression in Hs27 (**Figure 5H**) and BJ fibroblasts (**Figure S10F**). Flow cytometry also confirmed *SOX18* perturbation induced CD34 production but not PI16 (**Figure S10G**). Collectively, these results establish that our perturbative approach can elicit states *in vitro* that resemble states observed *in vivo*, and that both *KLF4* and *KLF5* can drive emergence of the universal fibroblast state.

### Causal drivers of fibroblast transcriptional states and cross-regulation between them

A natural question then arises of which of these candidate regulators is the dominant driver *in vivo*, as we observed homologous transcription factors could elicit similar phenotypes (**Figure 4A**). Our perturbation approach provides causal evidence that complements observational data. We compared the ability of each transcription factor to elicit universal or inflammatory gene programs *in vitro* to how correlated that transcription factor’s expression was with sensitive markers of those states within 96 fibroblast clusters in the Human Protein Atlas (**Figure 6A,B, Methods**). Outliers in this analysis categorized transcription factors as capable of modifying the phenotype *in vitro* (magenta), correlated *in vivo* (yellow), or both (blue). This analysis nominated *KLF4* and *OSR1* as causal drivers of the universal state *in vivo* out of 51 strongly correlated transcription factors. Notably, *Klf4* and *Osr2* are among the most differentially expressed transcription factors in mouse universal fibroblasts^53^ There was also a general enrichment of KLF transcription factors, with the activating factors *KLF2, KLF4,* and *KLF5* all promoting it *in vitro*, and the repressing factors *KLF3*, *KLF8*, and *KLF12* opposing it (**Figure 6A**)^91^. For the inflammatory state, the analysis intriguingly nominated *PLAG1* and *PLAGL1* as causal *in vivo* (**Figure 6B**). These ubiquitously expressed genes might be dismissed in purely correlative analyses, pointing to the interpretability advantages of our approach.

**Figure 6.**
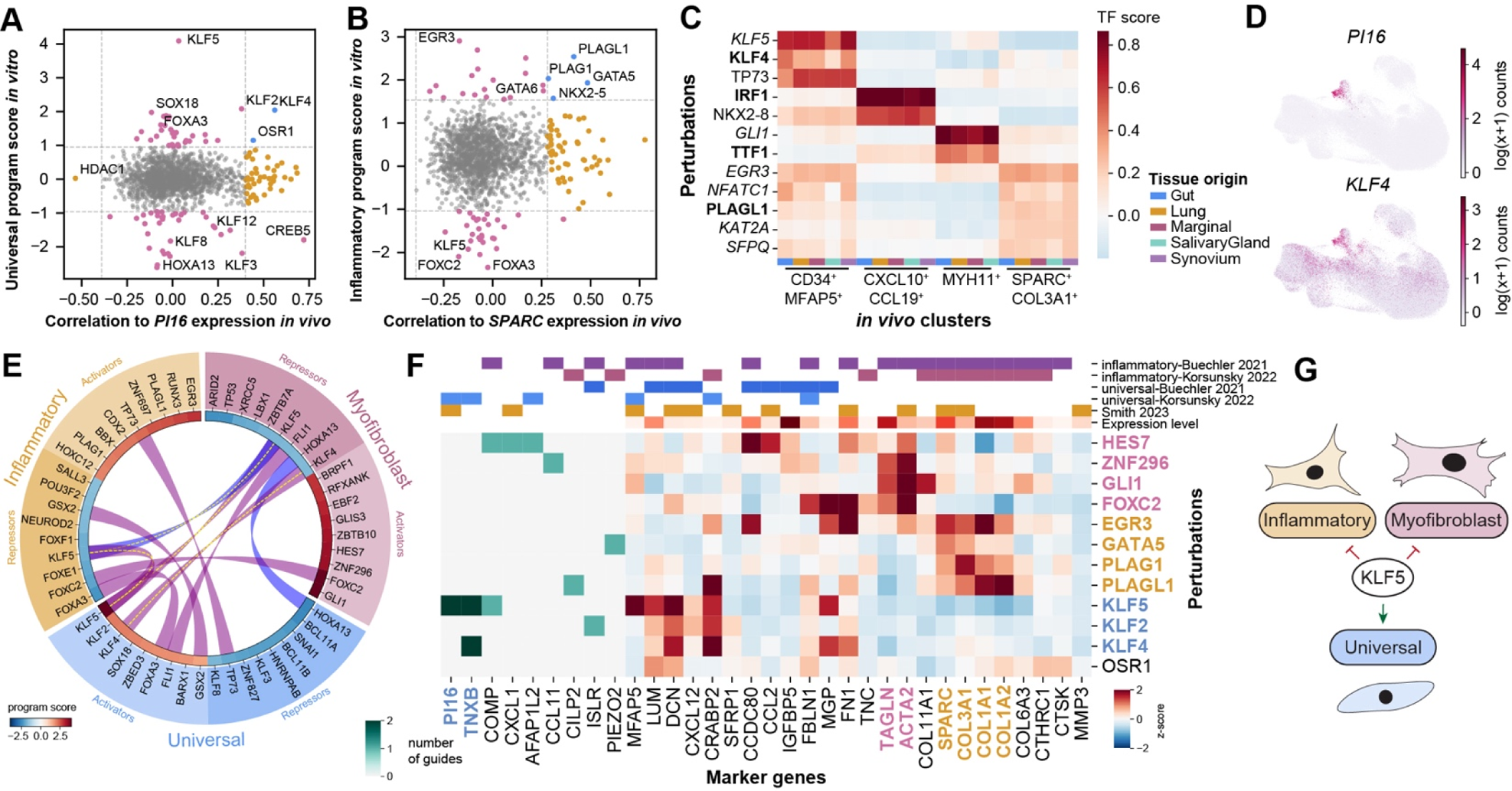
Causal drivers of fibroblast transcriptional states and cross-regulation between them. A. We compared the correlation of each transcription factor’s expression *in vivo* with PI16 expression in fibroblasts from the Human Protein Atlas2 vs. that transcription factor’s *in vitro* universal fibroblast program score. Yellow denotes highly correlated outliers (1.5 interquartile range, gray lines) in the *in vivo* data, magenta outliers in the *in vitro* program scores, and blue outliers in both analyses, which we infer to be likely causal drivers *in vivo*. B. Correlation of transcription factor expression *in vivo* with SPARC expression vs. the *in vitro* inflammatory program score, as described in A. C. We scored the similarity between the expression changes induced by each transcription factor in vitro vs. the differentially expressed genes within each in vivo cluster/tissue from an atlas of fibroblasts in inflammatory disease54. Plot includes the top two transcription factors for each cluster/tissue. Of these, the highest expressed transcription factor for each cluster is indicated in bold, and those expressed above the 50th percentile of all transcription factor expression are marked with italics. D. UMAP representation colored by expression of *PI16* (top) and *KLF4* (bottom) in fibroblasts collected from a large cohort of ovarian cancer patient samples profiled with scRNA-seq104. E. Cross-regulation between top activators and repressors of *in vivo*-associated gene programs. Arcs indicate co-occurrence of transcription factors across programs, with purple indicating opposing program effects, and blue conserved effects across programs. F. Heatmap summarizing the effects of top regulators from our analyses on key fibroblast marker genes. Inflammatory and universal markers from refs. 53, 54. General fibroblast markers: ref. 40 (see **Supplementary table 3**). Perturbations and genes are highlighted according to their related gene program. Magenta: myofibroblast transition, yellow: inflammatory program, blue: universal program. G. Schematic depicting the cross-regulation effect of *KLF5* in repressing (red) the inflammatory and myofibroblast cell states while promoting (green) the universal fibroblast state.

We observed similar results when we directly scored the similarity between transcription factor-induced phenotypes *in vitro* and transcriptional states observed *in vivo* in the stromal cell atlas of inflammatory disease (**Figure 6C**, Methods). *KLF4* and *KLF5* perturbations scored as most similar to the *CD34^+^MFAP5^+^*universal fibroblast state and *PLAGL1* and *EGR3* perturbations were hits for the *SPARC^+^COL3A1^+^* inflammatory fibroblast state. This analysis also supported a link between the *CXCL10*^+^*CXCL19*^+^ cluster and an *IRF1*-driven gene program (**Figure 4E**, **Figure S10A**). Finally, when we examined fibroblasts in a deeply sampled ovarian cancer dataset^84^, we observed a clear correlation between *KLF4* expression and *PI16* expression (**Figure 6D**). Overall, these analyses suggest *KLF4* is the primary cross-tissue driver of the universal fibroblast population *in vivo*.

Finally, throughout our analyses we observed a pattern in which activators of the universal fibroblast state were negative regulators of the inflammatory and myofibroblast states. This was apparent both among the top hit perturbations of our gene programs (**Figure 6E**, Methods), and in the effect of those hits on the expression of literature-defined markers for each of the states (**Figure 6F**, Methods). *KLF5* was the strongest hit in this regard (**Figure 6G**). This cross-regulation argues against a model in which *in vivo* fibroblast transcriptional states merely reflect different combinations of independent gene programs. Instead, states are interconnected and perhaps mutually exclusive. Inducing the universal state may therefore be a way to suppress disease-associated fibroblast phenotypes.

## Discussion

This study tested the ability to elicit *in vivo* transcriptional states *in vitro* using comprehensive perturbations of transcription factors via CRISPRa Perturb-seq. To enable exhaustive experiments targeting all transcription factors, we developed a multiplexing approach applicable to all Perturb-seq experiments based on overloading, achieving improved performance over past approaches by addressing a previously unappreciated saturation effect. We suspect that the competitive process causing differential capture of transcripts likely occurs in normal, singly-loaded droplets as well. Better understanding and correcting for these effects may therefore improve the overall sensitivity of single-cell RNA sequencing. The overloading approach is the natural choice whenever there is a high likelihood of interactions among perturbations, as effects of one strong perturbation may otherwise modify the effects of weaker ones measured concurrently. By pairing with approaches for installing programmed combinatorial perturbations^92,93^, it will instead enable scalable, directed profiling of genetic interactions.

These data yielded new insights into CRISPRa’s activity and specificity. We confirmed that guide efficacy is a significant challenge, with >50% of guides exhibiting no detectable on-target activity. A second, unexpected challenge was our identification of significant off-target effects driven by the seed region of the protospacer. Though present in past experiments, this effect was missed due to their limited phenotypic readout or smaller scale compared to the present study, where we have the resolution to see that they drive clustering and phenotypes of comparable magnitude to on-target effects. We suspect these off-target effects are an important consideration for all CRISPR effectors other than cutting, as they likely reflect transient activity of the attached domain as Cas9 scans the genome for target sites. The filtered, validated guide libraries developed here overcome these limitations, enabling compact future experiments targeting 1319 transcription factors using single guide RNAs (**Supplemental Table 4**). Combined with our multiplexing approach, screens targeting ∼1,000 transcription factors can be conducted in a single 10x HT lane, making it much easier to replicate our approach to examine transcriptional states in other cell types. Such screens could be used to nominate targeted lists of transcription factors for *in vivo* experiments^94–96^.

This work also establishes important advantages of CRISPRa as an overexpression tool. The most common concern with overexpression screens is the potential for neomorphic or nonphysiological phenotypes due to supraphysiological expression levels. In contrast to cDNA overexpression, the degree of activation by CRISPRa appears to be constrained by the endogenous regulatory environment: even the most potent guides activated unexpressed genes within the physiological expression range of transcription factors and activating already expressed genes yielded diminishing returns. The ability of CRISPRa to activate unexpressed genes was also a predictable product of epigenetic features of their promoters. Extending these predictions across the genome will help refine future library design. Other notable advantages of CRISPRa include the simplicity of multiplexing perturbations by co-expressing guides, the low cost of synthesis compared to cDNA libraries, the preservation of natural splicing and UTR regulation, and the ability to easily pair knockdown and activation^97^.

Our comprehensive, physiological CRISPRa perturbations of transcription factors in primary fibroblasts elicited expression programs mirroring four states observed *in vivo* (**Figure S10A**). Notably we employed no prior knowledge in the selection of our perturbations, such as human genetics or previous large-scale functional screens, which is valuable given that the origins and properties of these states are largely unknown. Our approach enabled causal insights that are impossible to obtain from purely observational data, highlighting the complementary nature of large-scale functional studies and cell atlas projects. We demonstrated co-regulation rather than co-expression among markers defining different states (**Figure 5A, 5C, 5F, 6F**), distinguished between causal and correlated transcription factors (**Figure 6A,B**), and observed cross-regulation between states (**Figure 6E,F**). Given our comprehensive perturbations, unelicited additional states may rely on specific *in vivo* conditions, reflect clustering artifacts, or result from combinatorial regulation. Nevertheless, our success in recapitulating states that recurred across multiple independent studies supports the idea that many transcriptional states reflect hard-coded regulatory logic that is conserved and elicitable *in vitro*.

Our most striking findings concern the universal fibroblast state. Critically, inducing this state can negatively regulate other fates, including the inflammatory state, which is expanded in various inflammatory diseases, and the myofibroblast state, which plays a central role in fibrotic disease. In addition to the putative multipotency and immunomodulatory properties^50^ of the universal state, this result underscores its importance as a therapeutic target. We identify *KLF4* as the dominant normal driver of the universal state *in vivo*, and we observe that other KLF transcription factors, including *KLF5*, strongly modify it *in vitro*. Previous studies have linked altered expression of *KLF2*^98–100^, *KLF4*^101–103^, and *KLF5*^83^ to fibrotic disease, which our results predict would accompany changes in the state composition of the fibroblast population. Importantly, our *in vitro* reconstitution approach now enables genome-wide experiments that would be impossible to perform *in vivo*. These experiments can identify synergistic inducers of the universal fibroblast state or repressors of the inflammatory fibroblast state, potentially leading to novel strategies targeting these states in disease.

Ultimately, our work demonstrates a systematic, generalizable, and scalable approach to dissect the regulatory logic of cellular transcriptional states, paving the way for more rational engineering of fibroblasts and other cell types.

## Supporting information

Supplemental Table 1 Composition of sgRNA libraries

Supplemental Table 2 Sparse gene expression programs

Supplemental Table 3 Curated set of fibroblast markers

Supplemental Table 4 Annotated guide activity metadata

Supplemental Table 5 Oligos List

Supplemental Table 6 Plasmids

## Acknowledgments

We gratefully acknowledge all the members of the Norman Lab for their valuable thoughts and discussion on this work. We thank Duaa H. Al-Rawi for her insight, and support and Ali Mohammad for his assistance with data transfer. We additionally thank the Shah lab (MSKCC) for facilitating analysis of the MSK-SPECTRUM fibroblasts. We acknowledge the use of the Epigenetics Research Innovation Lab, the Integrated Genomics Operation Core, and the and the Flow Cytometry Core Facility, which is funded by the NCI Cancer Center Support Grant (CCSG, P30 CA08748), Cycle for Survival, and the Marie-Josée and Henry R. Kravis Center for Molecular Oncology. This work was funded by NIH Director’s New Innovator Award DP2 GM140925, Damon Runyon Cancer Research Foundation Dale Frey Award AWD-GC-259296, and NIH U01 HG012103 (to T.M.N.). R.C.A. is a Robert Black Fellow of the Damon Runyon Cancer Research Foundation (DRG-2462-22) and D.D.O. was supported by NIH T32 GM132083. This research was funded in part through the NIH/NCI Cancer Center Support Grant P30 CA008748.

## Author contributions

Conceptualization, K.M.S., R.C.A., and T.M.N. Methodology, K.M.S., R.C.A., A.T., E.M., and T.M.N. Software, K.M.S., A.T., and T.M.N. Formal analysis, K.M.S., A.T., K.G., E.M., and T.M.N. Validation, R.C.A. Investigation, K.M.S., R.C.A., E.M., and D.D.O. Writing – Original Draft, K.M.S., R.C.A., A.T., and T.M.N. Writing – Review and Editing, K.M.S., R.C.A., A.T., and T.M.N. Visualization, K.M.S., R.C.A., A.T., K.G., E.M., and T.M.N. Supervision, T.M.N. Funding Acquisition, T.M.N. *The authors declare the following competing interests*: T.M.N. is an author on U.S. Patent No. 11,214,797B2, related to Perturb-seq. The authors otherwise declare no competing interests.

## Supplemental information

Document S1. Figures S1–S9 and Supplemental Figure legends

Table S1. Composition of sgRNA libraries, related to Figure 1, Figure 2

Table S2. Sparse gene expression programs for active perturbations, related to Figure 4

Table S3. Curated set of fibroblast state markers, related to Figure 5

Table S4. Annotated guide activity and metadata, related to Figure 2, Figure 4, Figure 6

Table S5. Oligos List, related to key resources table

Table S6. Plasmids, related to key resources table

## Methods

### CRISPR construct design/procurement

Components of VPH (vp64, p65, and HSF1) were ordered from IDT, PCR amplified and subcloned into pHR-UCOE-EF1a-Zim3-dCas9-P2A-mCherry, a gift from Marco Jost & Jonathan Weissman (Addgene plasmid # 188766; http://n2t.net/addgene:188766; RRID:Addgene_188766). This vector was further modified to have the SFFV promoter in place of EF-1α using the 2x NEBuilder® HiFi DNA Assembly Master Mix (NEB,E2621). Whole plasmid sequencing was performed by Plasmidsaurus using Oxford Nanopore Technology with custom analysis and annotation. pHR-UCOE-SFFV-VPH-dCas9-P2A-mCherry is designated pRCA422 and UCOE-SFFV-VPH-XTEN-dCas9-IRES2-BFP-T2A-Blast is designated pTMN140.

### Guide vectors

pRCA360 (GFP) and pRCA595 (mCherry) guide vectors were modified from pBA904, a gift from Jonathan Weissman (Addgene plasmid # 122238; http://n2t.net/addgene:122238; RRID:Addgene_122238). pJR107 was a gift from Marco Jost & Jonathan Weissman (Addgene plasmid # 187245; http://n2t.net/addgene:187245; RRID:Addgene 187245). For cloning of guide libraries or single guides, pRCA360, pRCA595, or pJR107 was sequentially digested using BstXI and BlpI (NEB) in their respective buffers (NEBuffer 3.1: B6003S, NEB CutSmart: B6004S) with gel purification to ensure efficiency. For single guide cloning, oligos were ordered with overhangs: Forward (5’-TTGN_20_GTTTAAGAGC-3’) and reverse (5’-TTAGCTCTTAAACN_20_ CAACAAG-3’) with reverse complemented guide sequence from IDT (see **Supplemental Table 5** Oligos List). 1µL of each F and R 100µM oligo is added to 48uL IDT Duplex buffer (IDT, 11-01-03-01) and heated to 95°C for 2 minutes before gradually cooled. Annealed oligos are diluted 1:20 with DNase free water and 1uL is used in a Hi-T4 (NEB, M2622S) ligation reaction at room temperature for 1 hour. Ligated products are transformed into NEB® Stable Competent E. coli (NEB, C3040H) as per manufacturer’s protocol and plated on Carbenicilin plates (Teknova, L1010) grown at 32°C overnight. Single colonies are picked, grown overnight at 32°C, miniprepped or midiprepped with QIAGEN kists (Qiagen, 27104 or 12945), and sequence checked by Plasmidsaurus.

### cDNA overexpression vectors

Lentiviral vectors overexpressing cDNA (TFORF0983 – KLF5, TFORF0985 – KLF4, TFORF3549 – GFP) were gifts from Feng Zhang. T2A-GFP was amplified using Q5® High-Fidelity DNA Polymerase (M0491, NEB) and plasmids were cut with EcoRI (NEB). Insert was cloned into these plasmids using the 2x NEBuilder® HiFi DNA Assembly Master Mix. A new empty vector construct was made using the newly generated TFORF Puro-T2A-GFP cut with NheI and MluI to remove the expressed cDNA. All vectors were sequenced checked by Plasmidsaurus

### Cell culture

hTERT RPE-1 Retinal Epithelium (RPE-1, ATCC CRL-4000), BJ fibroblasts (ATCC, CRL-2522), K562 (ATCC, CCL-243), and Hs27 foreskin fibroblasts (ATCC, CRL-1634) cells were purchased from American Type Culture Collection (ATCC). Lenti-X™ 293T Cell Line was purchased from Takara Bio (632180). hTERT RPE-1 were maintained in DMEM:F12 media (Gibco, 11320033) 10% Fetal Bovine Serum (FBS) (VWR, 97068-085), 100U penicillin and 100μg/mL streptomycin (Gibco, 15140122), and 0.01mg/mL Hygromycin B (Thermo Fisher, 10687-010). BJ and Hs27 fibroblasts are grown in DMEM (ATCC 30-2002) supplemented with 10% FBS. Lenti-X™ 293T is maintained in DMEM (Gibco, 11965092) with 10% FBS and Pen/Strep. Cells are passaged with CTS™ TrypLE™ Select Enzyme (Thermo Fisher, A1285901). K562 were maintained in RPMI-1640 (Gibco, 11875135) supplemented with 10% FBS and Pen/Strep.

### Virus packaging

Lentivirus was produced by co-transfecting LentiX™ 293T cells with transfer plasmids and standard packaging vectors (psPAX2, pMD2G) using TransIT-LTI Transfection Reagent (Mirus, MIR 2300). When required, virus supernatant was concentrated using the Lenti-X™ concentrator (Takara Bio, 631231).

### CRISPR cell line generation/procurement

RPE-1-ZIM3 CRISPRi cell line was obtained from Replogle and Saunders *et al*., 2022^62^. RPE-1-VPH cell line was generated by transducing RPE-1 from ATCC with 20X concentrated VPH lentivirus (pRCA422). mCherry+ cells were sorted into single cell clones in 96-well plate format. Single cell clones were grown and tested for uniform upregulation of CD55 (expressed) and CD45 (non-expressed) cell surface markers with a final clone selected for stable and homogenous target upregulation. BJ-VPH (pTMN140) and Hs27-VPH (pRCA422) cell lines were generated by sorting for either BFP-low or mCherry-low expression, respectively. CRISPRa activity of the sorted pool was confirmed by measuring cell surface expression of CD55 and CD45 as described below.

### Bulk characterization of cell lines

*RNA-seq*-RPE-1 cells were plated at a density of 300k per well into a 6 well plate. Cells were harvested 2 days later using RNEasy Plus Mini Kit (Qiagen, 74134) with QIAshredder (Qiagen, 79656) for homogenization and eluted in nuclease-free water. 500k Hs27 cells were plated on 100mm dishes. After 2 days, cells were harvested for RNA using the RNEasy Plus Mini Kit (Qiagen, 74134) and eluted with nuclease-free water. After RiboGreen quantification and quality control by Agilent BioAnalyzer, 500 ng of total RNA with RIN values of 10 underwent polyA selection and TruSeq library preparation according to instructions provided by Illumina (TruSeq Stranded mRNA LT Kit, catalog # RS-122-2102), with 8 cycles of PCR. Samples were barcoded and run on a NovaSeq 6000 in a PE100 run, using the NovaSeq 6000 S2 or S4 Reagent Kit (200 Cycles) (Illumina). An average of 50 million paired reads was generated per sample. Ribosomal reads represented 1.8-5.0% of the total reads generated and the percent of mRNA bases averaged 92%.

*ATAC*-50,000 freshly harvested RPE-1 and Hs27 cells were sent to MSKCC’s Epigenetics Research Innovation Lab for processing. ATAC was performed as previously described^105^ using the Tagment DNA TDE1 Enzyme (Illumina, #20034198). After quantification of the recovered DNA fragments, libraries were prepared using the ThruPLEX®DNA-Seq kit (R400676, Takara) following the manufacturer’s instructions, purified with SPRIselect magnetic beads (B23318, Beckman Coulter), quantified using a Qubit Flex fluorometer (ThermoFisher Scientific) and profiled using a TapeStation 2200 (Agilent). The libraries were sent to the MSKCC Integrated Genomics Operation core facility for sequencing on an Illumina NovaSeq 6000 (aiming for 50-60 million 100bp paired-end reads per library).

*CUT&RUN*-500,000 freshly harvested RPE-1 and Hs27 were sent to MSKCC’s Epigenetics Research Innovation Lab for processing. CUT&RUN was performed using the CUTANA™ ChIC/CUT&RUN Kit (Epicypher #14-1048) and the following antibodies: rabbit anti-H3K27me3 (CST, #9733S); rabbit anti-H3K27ac (Epicypher, #13-0059) rabbit anti-H3K4me3 (Epicypher, #13-0041); rabbit anti-mouse IgG (Epicypher, #13-0042). After quantification of the recovered DNA fragments, libraries were prepared using the ThruPLEX®DNA-Seq kit (R400676, Takara) following the manufacturer’s instructions, purified with SPRIselect magnetic beads (B23318, Beckman Coulter), quantified using a Qubit Flex fluorometer (ThermoFisher Scientific) and profiled using a TapeStation 2200 (Agilent). The libraries were sent to the MSKCC Integrated Genomics Operation core facility for sequencing on an Illumina NovaSeq 6000 (aiming for 5-10 million 100bp paired-end reads per library).

### Guide library design and cloning

*RPE-1-E150 (essentials)* – Guides for the small RPE-1 essentials library were selected towards 140 targets present within 13 functional clusters emerging from Replogle and Saunders *et al*., 2022^62^ RPE-1 essentials experiment (Supplemental Table 1). A total of 140 guides (RPE-1 essentials, Protospacer A for the 140 selected targets) and 16 non-targeting guides (sampled from Horlbeck *et al.,* 2016^55^ non-targeting controls) were selected. The guides were ordered as IDT oPool with overhangs for cloning into to the guide vector (pRCA360, overhangs: 5’-agtatcccttggagaaccaccttgttg(N_20_)-gtttaagagctaagctggaaacagcatagcaag-3’). To produce double stranded product for cloning, equimolar amounts of oligo pool and reverse primer (**Supplemental Table 5**: Oligo Library cloning R: cttgctatgctgtttccagc) were mixed with 2X KAPA HiFi master mix (Roche, 07958935001) and incubated for 10 min at 72°C. Following cleanup with a 5ug Monarch PCR& DNA clean up kit (T1030S), the library was cloned into digested pRCA360 using NEBuilder HiFi DNA Assembly (E2621S). A 5:1 insert to backbone HiFi mastermix reactions was performed for 1 hour at 50°C and purified using the Monarch® PCR & DNA Cleanup Kit (New England Biosciences, T1030). Electroporation was performed with an Eppendorf Eporator into Endura Electrocompetent Cells (LGC, 60242-2) according to manufacturer’s protocol. Overnight cultures are grown at 32°C for 18 hours before midiprep.

*CRISPRa-TFs* – We performed guide selection on the list of previously annotated and curated human transcription factors (MORF library^13^). We cross referenced all genes within the MORF library with the CRISPRa guide library described in Replogle, Bonnar, and Pogson *et al.*, 2022^56^(i.e. top 6 ranked from Horlbeck *et al*., 2016^55^). We further supplemented the guide list with the top 6 CRISPick^26,57^ (https://portals.broadinstitute.org/gppx/crispick/public) designs (CRISPRa, SpyoCas9) for each target. After strict filtering for off target binding, as described below, we ultimately selected 4 guides from Replogle, Bonnar, and Pogson *et al.*, 2022 and the top 2 CRISPick guides for the final library. Where there were not enough guides post filtration, or guides were unavailable for the specified target, additional guides from the other library were selected.

Filtration criteria was as follows: 1) candidate off-target sites for each protospacer were identified as i) having an NGG PAM and matching the 10 bp PAM-proximal sequence of the protospacer and ii) having less than 5 total mismatches. This search was limited to the accessible chromatin states within the Roadmap Epigenomics Project Core 15-state model of chromatin state^106^ within: NHDF-Ad Adult Dermal Fibroblast Primary Cells, i.e. all states except: 9-Heterochromatin, 13-Repressed PolyComb, 14-Weak Repressed PolyComb, 15-Quiescent/Low. 3) Each candidate site was assigned to the nearest gene and distance was calculated as absolute distance to the gene’s TSS. 3) A score for each site was calculated as: *log2(Distance*number of mismatches)*, where a lower score would suggest the guide more likely to be active at the site 4) a bimodal distribution of scores was observed, therefore a mixture model of two gaussians was fit and scipy.stats.norm.ppf(0.005) applied to the higher gaussian to threshold the distribution of high likely off-target binding from low likely off-target binding sites. Only sites that have a score less than the threshold were considered a “true off-target site”. Guides with more than 3 “true off-target sites” were excluded from the libraries. Additional filtration criteria included protospacers with three consecutive Ts (TTT) and those mapping to multiple sites in the genome (mostly attributed to paralog genes). If, after filtration and guide selection, the total number of guides per target from both libraries was less than 4; first, TTT filtered guides, then paralogs were recovered, where appropriate, to reach a total of 4 guides. Control guides were also strictly filtered to contain zero “true off-target sites” and at most one site that matches the 10 bp PAM-proximal sequence. Final transcription factor and control library compositions are described in **Supplemental Figure 1B** and full libraries available in **Supplemental Table 1.**

Libraries were ordered with overhangs for cloning (5’-agtatcccttggagaaccaccttgttg(N_20_)-gtttaagagctaagctggaaacagcatagcaag-3’) as an oligonucleotide pool (Twist Bioscences) and amplified using primers (Oligo Library cloning F: agtatcccttggagaaccacc, Oligo Library cloning R: cttgctatgctgtttccagc, see **Supplemental Table 5**) for 8 cycles with 2X KAPA HiFi master mix (Roche, 07958935001). Double stranded oligo cleanup and library cloning was performed as described above for the RPE-1 essentials library.

### Library Verification

Prior to virus generation libraries were sequenced to verify representation. Libraries were PCR amplified using 2X KAPA HiFi master mix (Roche, 07958935001) from purified midipreps with primers containing Illumina P5 and P7 adapters and Nextera i7 indices. (oJR232, oJR233, see **Supplemental Table 5** Oligos List). Following 2X cleanup with ProNex Size-Selective Beads (Promega, NG2001), sequencing was performed on Miseq with PE70 read structure. Resulting library representations are shown in **Supplemental Figure 1A,B.**

### RPE-1 essentials benchmarking experiment

Lentivirus was packaged for the RPE-1-E150 library as described above and the titer measured through infection of 250,000 RPE-1-ZIM3 with varying virus volumes. Infections were performed at the time of cell plating with 8 µg/ml polybrene. For the full-scale experiment 30M cells were infected across 6 T175s for a target MOI of 0.1. On day 3, an infection rate 14% was obtained and cells were sorted for high purity GPP+ (fluorescent marker for pRCA360 guide vector) by FACS (FACSAria III, and FacsSymphony S6, BD Biosciences). Post sort, cells were replated and cultured for additional 6 days. A timepoint of 6 days post infection was selected to minimize the dropout caused by targeting essential genes, while allowing sufficient time for phenotypic consequences to be captured within the experiment. The recommended protocol for adherent cell lines within “Single Cell Suspensions from Cultured Cell Lines for Single Cell RNA Sequencing” (10x Genomics Demonstrated Protocol, CG00054, Rev B) was followed to prepare cells for single-cell sequencing with a few deviations: 1) Two rounds of cell filtration through a 30 μm MACS SmartStrainers (Milenyi Biotec, 130-098-458) were performed after the initial cell resuspension. 2) a higher than recommended final cell concentration of 10,000 cells/μL was targeted for final resuspension 3) resuspension was performed in DPBS (Gibco, 14190144) containing 1% BSA (UltraPure BSA, 50 mg/mL, Ambion: AM2616) 4) Scienceware Flowmi Cell strainers 70 μm (H13680-0070) were used after final resuspension prior to cell counting. A cell concentration of ∼9700 cells/μL and >90% cell viability (Countess II, ThermoFisher) was obtained before loading. Cells were then encapsulated over 2 lanes using the Chromium Controller (10x Genomics) using the Chromium Next GEM Chip G Single Cell Kit (10X Genomics, 1000120) following the Chromium Single Cell 3’ Reagent Kits User Guide (v3.1 Chemistry Dual Index) with Feature Barcoding technology for CRISPR Screening (CG000316, Rev D) with the goal of recovering 125,000 cells per lane (prior to quality control and cell number filtering). Libraries were generated using the 3’ Feature Barcode Kit 16 rxns (10X Genomics, 1000262), Dual Index Kit TT Set A 96 rxns (10X Genomics, 1000215, and Dual Index Kit NT Set A 96 rxns (10X Genomics, 1000242) as per protocol. The following modifications were made to the cell loading protocol: 1) the expected cell recovery was empirically adjusted based on prior loading rate experiments for this cell line 2) BSA was added to the cell suspension/loading master mix for a final concentration of 1% BSA upon loading.

### RPE-1 TF-CRISPRa Experiment

Lentivirus was packaged for the TF-CRISPRa library as described above and the titer measured through infection of 250,000 RPE-1-VPH with varying virus volumes. Infections were performed at the time of cell plating with 8 ug/ml polybrene. For the full-scale experiment 30M cells were infected across 6 T175s for a target MOI of 0.05. After 2 days the infection rate was checked, obtaining a rate of 6%, and cells were sorted for high purity GPP+ (marker for pRCA360 guide vector) by FACS (FacsSymphony S6, BD Biosciences). Post sort, cells were replated and cultured for a total of 8 days post infection. The recommended protocol for adherent cell lines within “Single Cell Suspensions from Cultured Cell Lines for Single Cell RNA Sequencing” (10x Genomics Demonstrated Protocol, CG00054, Rev B) was followed to prepare cells for single-cell sequencing with a few deviations. 1) two rounds of cell filtration through a 30 μm MACs SmartStrainers (Milenyi Biotec, 130-098-458) were performed after the initial cell resuspension. 2) a higher than recommended final cell concentration of 10,000 cells/μL was targeted for final resuspension 3) resuspension was performed in DPBS (Gibco, 14190144) containing 1% BSA (UltraPure BSA, 50 mg/mL, Ambion: AM2616) 4) Scienceware Flowmi Cell strainers 70 μm (H13680-0070) were used after final resuspension prior to cell counting. A cell concentration of ∼10,000 cells/μL with >90% cell viability (Countess II, ThermoFisher) and >90% GFP+ (Attune, Invitrogen, ThermoFischer) was obtained before loading. Cells were then encapsulated over 10 lanes using the Chromium X (10x Genomics) using the Chromium Next GEM Single Cell 3’ HT Kit v3.1, 48 rxns (10X Genomics, 1000348) following the Chromium Next GEM Single Cell 3’ HT Reagent Kits v3.1 (Dual Index) with Feature Barcode technology for CRISPR Screening (CG000418, Rev D) with the goal of recovering 150,000 cells per lane (prior to quality control and cell number filtering). The following modifications were made to the cell loading protocol 1) the expected cell recovery was empirically adjusted based on prior loading rate experiments for this cell line 2) BSA was added to the cell suspension/loading master mix for a final concentration of 1% BSA upon loading.

### Hs27 TF-CRISPRa Experiment

Lentivirus was packaged for the TF-CRISPRa library as described above and the titer measured through infection of 1M Hs27-VPH cells with varying virus volumes. Infections were performed at the time of cell plating with 8 ug/ml polybrene. For the full-scale experiment 45M cells were infected across 15-150mm dishes for a target MOI of 0.1. After 2 days the infection rate was checked, obtaining a rate of 12.3%, and cells were sorted for high purity GPP+ (fluorescent marker for pRCA360 guide vector) using the SONY SH800 sorter. Post sort, cells were replated, cultured and expanded for a total of 12 days post infection. Prior to 10X loading, cells were sorted for GFP+ and measured to be >99% GFP+. The recommended protocol for adherent cell lines within “Single Cell Suspensions from Cultured Cell Lines for Single Cell RNA Sequencing” (10x Genomics Demonstrated Protocol, CG00054, Rev B) was followed to prepare cells for single-cell sequencing with a few modifications: 1) Two rounds of cell filtration through a 100 μm MACs SmartStrainers (Milenyi Biotec, 130-098-458) were performed after the initial cell resuspension. 2) a higher than recommended final cell concentration of 10,000 cells/μL was targeted for final resuspension 3) resuspension was performed in DPBS (Gibco, 14190144) containing 1% BSA (UltraPure BSA, 50 mg/mL, Ambion: AM2616) 4) Scienceware Flowmi Cell strainers 70 μm (H13680-0070) were used after final resuspension prior to cell counting. A cell concentration of ∼10,000 cells/μL with >99% GFP+ (Attune, Invitrogen, ThermoFisher) and >90% cell viability (Countess II, ThermoFisher) was obtained before loading. Cells were then encapsulated over 10 lanes using the Chromium X (10x Genomics) following the Chromium Next GEM Single Cell 3’ HT Reagent Kits v3.1 (Dual Index) with Feature Barcode technology for CRISPR Screening (CG000418, Rev D) with the goal of recovering 100,000 cells per lane (prior to quality control and cell number filtering). The following modifications were made to the cell loading protocol 1) the expected cell recovery was empirically adjusted based on prior loading rate experiments for this cell line 2) BSA was added to the cell suspension/loading master mix for a final concentration of 1% BSA upon loading.

### Perturb-seq Library Preparation and Sequencing

Sample processing and library preparation was generally performed as recommended within 10x Genomics Chromium Single Cell 3’ Reagent Kits User Guide (v3.1 Chemistry Dual Index) with Feature Barcoding technology for CRISPR Screening (CG000316, Rev D) (RPE-1-Essentials) or 10x Chromium Next GEM Single Cell 3’ HT Reagent Kits v3.1 (Dual Index) with Feature Barcode technology for CRISPR Screening (CG000418, Rev D) (HT kit, TF-CRISPRa Library experiments) with the following modifications 1) During the Dynabeads cleanup step (step 2.1c, standard kit, step 2.1c HT kit), 2 times the recommended amount of Dynabeads MyOne SILANE was added to the cleanup mix and Nuclease-free Water was omitted (RPE-1 benchmarking), alternatively Nuclease-free Water was simply replaced with Dynabeads MyOne SILANE (TF-CRISPRa Library experiments). 2) The amount of cDNA amplifications cycles (step 2.2d standard, 2.2d HT) was reduced 1-2 cycles from the lowest recommendation. These steps were found to reduce Dynabead clumping and increase the cDNA yield with equivalent input and cDNA cycles, and to produce sufficient material for library generation without overamplification. For library generation from the HT kit A and B libraries (i.e. the first or second pipette from the GEM well. Steps 1.4 e/h-row 3A and i/k-row 3B) were maintained separately throughout the protocol; libraries were not combined at Step 2.3A-ix but instead carried through individually. This was done to enable library level batch effect corrections in subsequent analysis. In the standard protocol, recombination is suggested after separate GEM-RT, cleanup, and cDNA amplification steps, introducing batch effects that are obscured upon library recombination. However, due to droplet settling, uneven cell numbers are observed across the A and B libraries the libraries (first and second pipette). To account for this, lanes were initially sequenced at low depth and re-sequenced to balance reads per cell after initial cell calling. For all Feature Barcode (CRISPR guide capture) library preps, all material, i.e. ∼150μL of guide containing supernatant, was retained during cDNA cleanup (Step 2.3d) and purified at a 1.2X SPRIselect ratio to the measured volume in step 2.3B-i. As done for the GEX libraries, Feature Barcode libraries were maintained separately and not recombined as specified in 4.1m. Instead, libraries were eluted in 50.5μL in step 4.1j and 50μL of each library was transferred to a new tube (4.1m).

After PicoGreen quantification and quality control by Agilent TapeStation, libraries were pooled equimolar and run on a NovaSeq 6000 or X in a PE28/88 run, using the NovaSeq 6000 S4 200-cycle or X 10B or 25B 100-cycle Reagent Kit (Illumina). The loading concentration was 0.6-1.0 nM and a 1% spike-in of PhiX was added to the run to increase diversity and for quality control purposes. The runs yielded on average 326M reads per sample.

### Profiling of genome-wide dCas9 binding by CUT&RUN, related to Figure 3F

*Generation of dCas9 construct and cell lines for CUT&RUN*-A dCas9-rabbit Fc domain fusion protein (vector pEM022) was constructed by adding a linker sequence and the rabbit Fc domain to the C-terminus of dCas9 within a lentiviral vector (pEM021, originating from pRCA422). The rabbit Fc sequence and a 5’ linker (obtained from *Wang et al*.^107^) was codon-optimized and ordered from IDT as a gBlock, which was cloned via HiFi Assembly into a NotI digest of this intermediate vector. Lentivirus was produced by co-transfecting LentiX 293T cells with pEM022, psPAX2, and pMD2G, and pooled lentiviral stocks harvested at 48 hr and 72 hr post-transfection were concentrated 20x with the Lenti-X Concentrator kit (Takara #631232). K562 cells (ATCC, CCL-243), either wild-type or harboring a lentiviral integrated eGFP driven by the SV40 promoter, were transduced with 500 µL concentrated virus per million cells by spinfection in the presence of 8 µg/mL polybrene and sorted for BFP positivity after 48 hr to establish stable dCas9-Fc cell lines. This cell line was similarly transduced with lentivirus encoding individual sgRNAs produced in arrayed format (vector pRCA595 or pRCA360). At 48 hr post-transduction, sgRNA-containing cells were selected with puromycin (1.5 µg/mL media) for 4 days.

*Guide dependent dCas9 CUT&RUN-* profiling was performed on each of these cell lines (no sgRNA and each of 6 sgRNAs with different seeds, **Supplementary Table 5**) using the CUTANA ChIC/CUT&RUN Kit Version 4 (EpiCypher #14-1048). 1.5×10^6^ cells of each cell line were harvested and divided among three technical replicates of 5×10^5^ cells. Before each experiment, viability was confirmed >95% by Trypan Blue (Countess II FL, Life Technologies) to reduce background signal from dead cells. Since the dCas9-Fc fusion protein obviates the need for a primary antibody, the protocol was performed with slight modifications: Section IV was skipped, and pAG-MNase binding was started at Step V.32 with cells still in Antibody Buffer (which contains EDTA, chelating Ca^2+^ to prevent nonspecific pAG-MNase cleavage before the digestion step). For pAG binding to dCas9-Fc (step V.33), reactions were incubated for 2 hr on a nutator at 4°C. After incubation, cells were washed three times with Cell Permeabilization buffer, and the protocol was continued without modifications. Sequencing libraries were prepared using the CUTANA CUT&RUN Library Prep Kit (EpiCypher #14-1001) with no modifications. Individual libraries were sequenced on an Illumina NovaSeq X to a target depth of 10 million PE75 reads.

### CD55/CD45 cell surface marker staining, related to Supplemental Figure 1B,C

100k BJ-VPH, 150k Hs27-VPH, or 300k RPE-1-VPH cells were transduced with lentiviruses of pRCA360 expressing sgRNA against CD55, CD45, or a non-targeting guide (NTC) with 8µg/mL polybrene in a 6-well plate. Puromycin (Gibco, A1113803) selection (1.2µg/mL) was done at day 2 for 4 days for fibroblasts but not RPE-1. On day 8, cells were detached using TrypLE and washed once with BD Pharmingen™ Stain Buffer (BD, 554656) + 2mM EDTA (Thermo Fisher, 15575020). Cells were stained with CD55-APC (BD, 555696) and/or CD45-PerCP-Cy5.5 (BD, 564105) for 30 minutes on ice in the dark, washed with stain buffer, and analyzed on BD LSRFortessa or Thermo Fisher Attune flow cytometer. Parental cell lines expressing GFP, BFP, or mCherry was used as single-color compensation controls. CD55-APC (BD, 555696) and CD81-PerCP-Cy5.5 (BD, 565430) was used as a single-color control. Flow cytometry data was analyzed using Flowjo software.

### PI16/CD34 cell surface marker stainingfor CRISPRa, related to Figure 5H, Supplemental Figure 10E, G

150k Hs27-VPH cells were transduced with lentiviruses of pRCA360 expressing sgRNA against KLF5, KLF4, ASCL5, SOX18, or a non-targeting guide (NTC) with 8µg/mL polybrene. On day 3, puromycin selection (1.2µg/mL) was done for 2 days on cells lifted with TrypLE. Cells were cultured and expanded with growth media until harvest at day 13 or 17. For this assay, cells were detached using Non-Enzymatic Cell Dissociation Solution (ATCC, 30-2103) and washed with stain buffer + 2mM EDTA. Cells were stained with PI16-APC (Miltenyi, 130-111-177) and/or CD34-PerCP-Cy5.5 (BD, 347213) for 30 minutes on ice in the dark, washed, and analyzed on BD LSRFortessa or Thermo Fisher Attune flow cytometer. Parental cell lines expressing GFP, BFP, or mCherry was used as single-color compensation controls. CD55-APC (BD, 555696) and CD81-PerCP-Cy5.5 (BD, 565430) was used as a single-color control. Flow cytometry data was analyzed using Flowjo software.

### PI16/CD34 cell surface marker staining for cDNA overexpression, related to Figure 5H, Supplemental Figure 10F

150k Hs27 wiltdype or 100k BJ wildtype cells were transduced with lentiviruses of TFORF Puro-T2A-GFP or TF-ORF Puro-T2A-BFP expressing cDNA of KLF5, KLF4, or empty vector (EV) with 8µg/mL polybrene. On day 2, puromycin selection (1.2µg/mL) was done for 2 days on cells lifted with TrypLE. Cells were cultured and expanded with growth media until harvest at day 13. For this assay, cells were detached using Non-Enzymatic Cell Dissociation Solution (ATCC, 30-2103) and washed with stain buffer + 2mM EDTA. Cells were stained with PI16-APC (Miltenyi, 130-111-177) and/or CD34-PerCP-Cy5.5 (BD, 347213) for 30 minutes on ice in the dark, washed, and analyzed on BD LSRFortessa or Thermo Fisher Attune flow cytometer. Parental cell lines expressing GFP was used as single-color compensation controls. CD55-APC (BD, 555696) and CD81-PerCP-Cy5.5 (BD, 565430) was used as a single-color control. Flow cytometry data was analyzed using Flowjo software.

### ACTA2 immunofluorescence, related to Figure 5B, Supplemental Figure 10B

Buffer composition are as follows: 1) Wash buffer: PBS (Thermo Fisher, 14040117) + 0.05% Tween-20 (Sigma Aldrich, P7949), 2) Blocking buffer: wash buffer with 5% BSA (Thermo Fisher, J6509722). Hs27-VPH were transduced with lentiviruses of pRCA360 expressing sgRNA against HES7, ZNF296 or a non-targeting guide (NTC) with polybrene. On day 2, puromycin selection (1.2µg/mL) was done for 2 days on cells lifted with TrypLE and replated. Cells were cultured and expanded with growth media until day 17, where cells were detached, counted, and 2.5k cells were plated into 96-well plates optical plate (Cellvis, P96-1.5P). On day 18, cells were fixed with ice-cold methanol for 15 minutes at −20°C. Cells were blocked in blocking buffer for 1 hour at room temperature. ACTA2 antibody (Sigma Aldrich, A2547) diluted 1:2000 in blocking buffer was added to the plate and incubated overnight in the dark at 4°C. The next day, cells were washed for three times with wash buffer for 10 minutes each. Goat anti-mouse AF674 secondary antibody (Invitrogen, A-21236) was diluted 1:2000 in blocking buffer and added to plate for 1 hour at RT. Post-secondary antibody, the plate was washed three times with wash buffer for 10 minutes each. A 1:5000 solution of DAPI (4’,6-diamidino-2-phenylindole) (Thermo Fisher, 62248) diluted blocking buffer was added for 15 mins at room temperature to stain for DNA. Cells were then washed for three times with wash buffer before being imaged on a Nikon Eclipse Ts2. Images were analyzed using Fiji.

### Computational

#### Dataset processing and guide calling

cellranger-7.1.0 (10x Genomics) was used for alignment and UMI collapse of reads from gene expression (GEX) libraries to the transcriptome (reference: 10x Genomics GRCh38 version 2020-A). sgRNA reads were aligned to the protospacer library utilizing the cellranger count `--feature-ref` option with specified library inputs. Resulting filtered cell calls (i.e. droplets) were used for further cell assignment with droplet overloading. For the RPE-1 Essentials Library, cellranger aggr was run without downsampling `(--normalize none`) to aggregate the lanes and perform final guide assignment per droplet (Cellranger’s CRISPR Guide Capture Algorithm). Integrated libraries were imported with scanpy (version 1.7.2) and a final threshold of 5 UMIs was added prior to final cell assignment to account for failed protospacer thresholding (i.e. very low UMI threshold in cellranger output).

For TF-CRISPRa Library experiments, we used a 5 UMI cutoff to make guide assignments per droplet. This was effective given the great diversity of protospacers within the library leading to low background UMIs per protospacer. Libraries from TF experiments were aggregated with scanpy without read downsampling.

### Dataset filtering

*Quality filtering of droplets*. As discussed in the main text, total UMI count scaled with the number of cells loaded into each droplet. We removed a small number of droplets that had low library sizes for their occupancy level. Specifically, for each dataset we computed histograms of the log_2_ total UMI count within each droplet and then set thresholds based on visual inspection of the lower tail of the distributions. These thresholds depend on sequencing depth and are dataset-dependent:

| **Droplet occupancy** | **RPE-1 essentials** | **Hs27 CRISPRa, gemgroup ≤ 7** | **Hs27 CRISPRa, gemgroup > 7** | **RPE-1 CRISPRa** |
| --- | --- | --- | --- | --- |
| **1** | $2^{10.7}$ | $2^{11}$ | $2^{10.7}$ | $2^{11}$ |
| **2** | $2^{11}$ | $2^{11.5}$ | $2^{11}$ | $2^{11}$ |
| **3** | $2^{12}$ | $2^{11.7}$ | $2^{11.3}$ | $2^{11.3}$ |
| **4** | $2^{12.7}$ | $2^{12}$ | $2^{11.5}$ | $2^{11.7}$ |
| **5** | $2^{13.5}$ | | | |
| **6** | $2^{14}$ | | | |

We also applied dataset-dependent filters on mitochondrial reads to remove droplets that produced low complexity libraries. To set these filters we computed the total fraction of mitochondrial RNA reads, plotted histograms of total UMI count vs. fraction of mitochondrial reads to identify outliers, and discarded any droplets with fractions greater than the following thresholds: 0.075 for RPE-1 essentials, 0.05 for Hs27 CRISPRa, and 0.03 for RPE-1 CRISPRa.

Finally, as noted in the table above, we only kept droplets containing up to 6 cells for the RPE-1 essentials experiment, and up to 4 cells in the CRISPRa experiments.

*Filtering of control guides*. Our experiments incorporated numerous non-targeting control guides, some of which may inadvertently have targeting effects. To investigate, for each experiment we constructed an expression matrix consisting of all genes with mean expression greater than 1 UMI per cell (as columns) and all droplets containing only a single candidate control guide (as rows). We *z*-normalized the columns of this matrix and then computed pseudobulk expression profiles for each control guide by grouping by the control guide received and computing the mean. We then visually inspected a clustermap of these pseudobulk profiles to look for obvious outliers. We also computed a score by computing the L1 norm of the pseudobulk profiles, as outliers will have large norm due to the *z*-normalization. We made conservative, dataset-specific choices about thresholds. Candidate control guides we discarded are annotated in **Supplemental Table 4**, *off-target guides*.

### Identification of stably captured genes and normalization procedure

*Overview of procedure*. As described in the main text, our normalization and regression approach consists of three broad steps:

1. Identification of genes that are “stably” captured across different droplet occupancies, which are used to estimate a global scale factor that we apply to make droplets of different occupancies comparable to each other.
2. Identification of gene-specific scale factors that are applied to compensate for the variable capture efficiency we observe for different transcripts as droplets are overloaded.
3. Application of these scale factors, and then normalization relative to droplets that contain single control cells.

*Step 1: Global scale factors derived from stably captured genes*. We first computed global scale factors to adjust for differences in sequencing depth across droplets. This variability in total library size is a feature of droplet single-cell RNA sequencing workflows and is a larger source of heterogeneity than any of the perturbations we apply. As discussed in the main text, we also observed a saturation effect that appeared to affect different transcripts in different ways, with some becoming more likely to be observed in droplets of higher occupancy levels and others less likely. This likely indicates that some sort of competitive process takes place during library capture or amplification. We therefore derived a procedure to estimate library size that is independent of these effects.

We computed an expression matrix consisting of genes with mean greater than 0.5 UMI per cell (as columns) and all quality filtered droplets as rows. We normalize each row by the total UMI count within that droplet: the *j*th entry in the *i*th row is then the fraction of total UMIs captured in the *i*th droplet that correspond to gene *j*. Below we will refer to this matrix as *P*, as the rows can be viewed as probability distributions on expression. If we average together the rows of *P* corresponding to droplets containing the same number of cells, we thus get estimates of the average capture rate of that transcript according to droplet occupancy as visualized in **Figure 1E**.

We then wished to identify transcripts that were least affected by these shifts in capture efficiency, which we termed “stably captured.” Our notion was that these genes should have similarly shaped distributions (when normalized as in *P*) regardless of droplet occupancy. To quantify differences in the shapes of distributions, we used the Wasserstein distance. Specifically, for each gene we computed its distribution (across rows in *P*) for each droplet occupancy level, computed all possible pairwise comparisons among these distributions with the Wasserstein distance, and then summed the result to produce a “cost” for that gene. We performed this procedure independently for each gemgroup in case there were gemgroup-specific differences in capture efficiency and summed them to produce a “total cost.” Genes whose distributions vary little according to droplet occupancy (or gemgroup) will have a low total cost. The total cost did scale generally with the mean expression level of the gene (due to properties of Wasserstein distance), so we then computed a regression of log_10_ (total cost) as a function of log_10_ mean expression level) using Huber robust regression and subtracted the regression fit (i.e. computed the residual) to produce a “mean-adjusted total cost” for each gene.

We observed that some cell cycle regulators had low total cost but would serve as poor references for normalization due to their substantial single-cell heterogeneity. We therefore computed a measure of “excess CV” to identify particularly noisy genes. Specifically, for each gene we computed its coefficient of variation (CV). We then ordered genes by mean expression level and fit a nonlinear curve using a median filter (specifically, `medfilt(gene_cvs_mean_ordered, kernel_size=15)`). We then fit an interpolation to this curve using interp1d that allowed us to produce for each gene an estimate of the expected (median) CV given its observed mean expression level. The ratio of observed CV to expected CV was the “excess CV.” More variable genes will thus have excess CV > 1 (i.e. noisier than expected for their expression level).

We then identified stably captured genes using two thresholds: (1) the gene had to be in the bottom 30% of the mean-adjusted total cost distribution, and (2) the gene had to have excess CV < 1. We wanted to use these stably captured genes to estimate global scale factors for each droplet. Our idea was for each droplet we could infer an estimate of what its total library size would be if all genes were stably captured (i.e. not affected by the variability in capture rate discussed above). Our approach was to first fit a scaling constant for each droplet across stable genes:

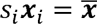

where *x_i_* is a vector composed of the (unnormalized) expression levels of stably captured genes in a given droplet, 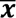 is the mean raw expression of stably captured genes across all droplets with the same occupancy level, and *s_i_* is a scale factor fit by Huber robust regression (to prevent undue influence of outliers or stochasticity). Put simply, this procedure identifies the single number that best scales the UMI counts of stably captured genes observed in droplet *i* to be proportional to those observed in the average droplet of the same occupancy level.

Finally, we then used these coefficients to produce estimates of total library size (rather than just the scale of stably captured genes). For each droplet, we multiplied the observed total UMI count *U_i_* by the fitted coefficient to produce an estimate of what the total UMI count would be if all genes behaved like stably captured genes. To enable comparisons across droplets of different occupancy levels, we then further normalized these by the median estimated UMI count across all droplets of the same occupancy level to produce our final scale factors:

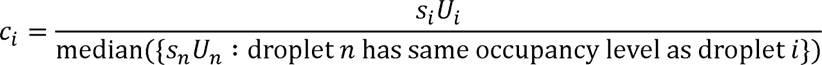

**Figure 1F** shows the distributions of *s_i_U_i_* for droplets of different occupancy levels in the RPE-1 essentials experiment (without the median normalization in the denominator). Note that they are much better resolved by droplet occupancy level than the raw total UMI counts, despite the procedure not taking droplet occupancy into account during fitting.

Overall, this procedure produces an estimate of total library size *C_i_* that ranges roughly between 0.75 and 1.25 that can be thought of as an estimate of that droplet’s library size—a droplet with a library size of 0.75 for example is undersequenced relative to other droplets and has only 75% of the expected total UMI count. It thus preserves some of the variation in total library size that would be erased by normalization procedures that for example force all libraries to have a fixed total count. Moreover, because this measure is computed based on stably captured genes, it can be used to normalize droplets of different occupancies to comparable levels.

*Step 2: Computing gene-specific scale factors to adjust for varying capture efficiency*. The previous procedure provides a global estimate of library size/sequencing depth for each droplet that is robust to the variability in transcript capture efficiency that we observed. We next wanted to develop first order corrections for this variability as well. To do this, we first computed a matrix 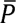 from *P* by (1) grouping together rows with the same occupancy levels and (2) for each group, calculating a 2%-trimmed mean for each gene. (We used trimmed means to try to make the procedure somewhat resilient to the stochastic variability in expression that is a feature of single-cell RNA sequencing.) The rows of 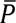 are then estimates of the average capture rate of all transcripts as visualized in Figure 1E.

To develop correction factors, we then computed a mean capture rate vector *µ* by computing the same trimmed mean calculation over all rows of *P* (independent of occupancy level). Our correction factors were then defined row-wise via:

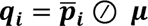

where ***q_i_*** and 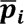 are the *i*th rows of matrices *Q* and 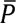 and 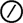 represents Hadamard (element-wise) division. The matrix *Q* contains our gene-level correction factors that we can apply to each droplet. Dividing the expression of gene *j* by *Q_ij_* in droplets containing *i* cells will adjust expression so that, on average, transcripts are captured at the same rate regardless of droplet occupancy level. In other words, it will “divide out” the variable capture rates seen in **Figure 1E**, on average.

Note: Gene-level factors were fitted independently within each gemgroup in case of gemgroup-specific differences due to library preparation or overall sequencing depth.

*Step 3: Final normalization.* We finally combine these ingredients to produce the final normalization. We proceed as follows:

A. We compute the matrix *P* of normalized expression for each droplet.
B. For each droplet *i*, we use the procedure of Step 1 above to compute a global library size estimate *C_i_*.
C. We use *P* and the procedure in Step 2 above to compute gene-level scaling factors *Q*.
D. We then apply these factors to adjust for variability in library size and transcript capture rate. If droplet *i* contains *k* cells, we normalize expression of gene *j* by 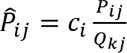, where 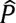 is the corrected normalized expression matrix.
E. Finally, we convert these estimates to deviations relative to expression seen in control cells within the same gemgroup. As described in our past work^20,62^ this procedure helps adjust for batch effects that arise during library prep. We accomplish this by “z-normalizing” relative to statistics derived from control cells (within droplets that contain only one cell). If gene *j* has mean *μ_j_* and standard deviation *σ_j_* in control cells, our final normalized expression estimate in droplet *i* is

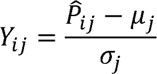

The advantage of this scaling of gene expression is that (1) droplets containing single control cells will have 0 mean expression on average for all genes and (2) there is a natural interpretation of the scale of coefficients in terms of “standard deviations of change relative to control cells.” For example, a score of 2 would indicate that a cell is two standard deviations above mean expression in control cells for a given gene.

### Computing mean guide phenotypes using linear regression

The normalization procedure described in the previous section places droplets of all occupancies on an equivalent expression scale and further represents our gene expression measurements in terms of deviations relative to expression in control cells. As a result, the task of linear regression is relatively simple. We construct a binary design matrix *X* (scaled so that each row sums to 1) with entries

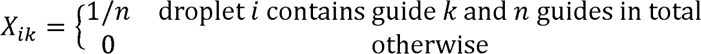

(Note: All control guides are treated as having the same “perturbation” which is called “control”.) We then perform the multiple linear regression

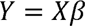

using scipy’s `lstsq` function. Coefficient *β_kj_* then corresponds to the inferred effect of guide *k* on gene *j*. We attach *p*-values to coefficients using the standard *t*-test. (Our specific implementation is based on the one in the pingouin library.) Because of the relative normalization, significant (non-zero) coefficients mark gene expression changes relative to (singlet) control cells. We produce *p*-values adjusted for multiple hypothesis testing using the Benjamini-Hochberg procedure.

In the RPE-1 essentials experiment we fit the model to all genes with mean greater than 1 UMI per droplet (and over all droplets containing up to 6 cells). In the Hs27 and RPE-1 CRISPRa experiments we fit the model to all genes with mean greater than 0.2 UMI per droplet (and over all droplets containing up to 4 cells).

### Benchmarking regression approach, related to Figure 1 and Supplemental Figure 2

*Baseline normalization used for comparison.* To provide a baseline gene expression normalization for comparison, we used the gemgroup z-normalization from our past work^20,62^. Specifically, we:

1. Normalize all droplets to have equal total UMI counts
2. Perform the “*z*-normalization relative to control cells” for each gene described in the previous section. The control cell means and standard deviations are determined independently within each gemgroup due to the batch effects we have described previously^62^.

The primary difference is thus that all droplets are equalized to have the same total UMI count, meaning the UMI saturation effect we describe in the main text is not accounted for. The same design matrix is used with both gene expression normalization approaches.

*Figure 1G-I*: We first tested reconstruction overall reconstruction reproducibility based on the inferred coefficients in *β*. Because Perturb-seq is typically conducted with relatively modest sampling per perturbation, some variability is expected. Our baseline comparison therefore consisted of taking all of the singlet droplets in the experiment, splitting them into two halves (held out vs. test), computing regressions within each split to produce *β*_held out_ and *β*_test_, and then comparing the results. Our metric was the Pearson correlation coefficient computed between *β*_held out_ and *β*_test_, each stacked into vector form. We then systematically downsampled the size of the test set, recomputed *β*_test_, and measured performance again. Across many downsamplings, this produced the calibration curve seen in in **Figure 1G**. We then fit an interpolation to this curve using interp1d, which allows us to map an observed correlation between *β*_held out_ and *β*_test_ to an equivalent number of singlet droplets in the test set. Finally, we then selected test sets consisting of equal numbers of cells contained in either singlet, doublet, triplet, or quadruplet droplets. (I.e. 16800 singlets vs. 8400 doublets vs. 5600 triplets vs. 4200 quadruplets.) We computed the corresponding *β*_test_, computed performance relative to the held-out set, used the calibration curve to map the observed performance to an equivalent number of singlet droplets, and then finally divided this number by the actual number of droplets in the test set. A performance of 1 by this metric then means that the given droplets perform equivalently to singlets: e.g. 8400 doublets yielded approximately the same performance as 8400 singlets, and so received a score of ∼1 in **Figure 1H**. That figure compares the performance of these regressions using the baseline normalization vs. our stably captured gene normalization.

**Figure S3B** explores an alternative comparison along similar lines. In this case the test set consisted of only of doublets, and we varied the number between 8400 (0.5x doublets) and 16800 (1x doublets), showing that 16800 doublets perform roughly equivalently to 16800 singlets in terms of reconstruction reproducibility. We next performed reconstructions using all droplets in the experiment containing up to 6 cells. We downsampled them to varying total numbers (**Figure S3C**) and compared reconstructions again to a held out set of 16800 singlets. **Figure 1I** shows the raw Pearson correlation coefficient between *β*_held out_ and *β*_test_ for these different choices of the test set (blue dots), with the magenta dots showing performance using the gemgroup z-normalization for comparison. The gray lines show the performance of the 16800 singlet test set that we use as a baseline comparison, again showing roughly equivalent performance on a droplet-for-droplet basis regardless of occupancy level.

Finally, **Figure S3D** compares heatmaps of the *β* coefficients computed using either all singlets in the experiment (33681) or all droplets in the experiment containing up to 6 cells, using either our stably captured genes-based normalization or the baseline gemgroup *z*-normalization. Inspection of the heatmaps of the latter reveals regions of structural differences driven by the UMI saturation effect that are absent in the former, which are largely visually indistinguishable.

*Figure 1J*: The previous comparisons were all based on the raw values of the inferred *β* coefficients. However, capturing these absolute effects may not be necessary for applications that require only accurate reconstruction of the relationships between perturbations. For example, clustering relies on pairwise comparisons and can be used to assign function to perturbations. We thus conducted parallel analyses to the previous ones using a different metric which we refer to as cophenetic correlation. Specifically, we compute *β*_held out_ and *β*_test_ as above, then compute the perturbation-perturbation correlation matrices *R*_held out_ and *R*_test_, and finally compute the similarity between these, again by stacking them and computing the Pearson correlation. Figure 1J shows the result of this for the experimental design in **Figure S3C**, showing that it is in general easier to reconstruct correlations than absolute effects, but that our stably captured gene normalization still improves performance. Accordingly, **Figure S3E** shows an equivalent analysis to **Figure 1G-H** using cophenetic correlation.

*Figure S2F-I*: Finally, we conducted equivalent analyses for the *p*-values assigned by our regression procedure to the coefficients in *β*. Due to the large-scale differences in *p*-values, we compared them using Spearman correlation instead of Pearson correlation. **Figure S3F** shows an analysis parallel to **Figure 1G-H**, showing that the gemgroup *z*-normalization severely distorts significance testing, which is substantially improved by the stably captured gene normalization. (Compare with **Figure S3D**, showing structural differences in the coefficients that are likely driving this divergence.) Figure S2G follows the design of Figure S2C, but for *p*-values. The relatively weak agreement here is driven partly by the low number of cells per perturbation, as we see similarly modest reproducibility using two samples of 16800 singlets (gray lines). An alternative comparison is to look at the number of differentially expressed genes (adjusted *p <* 0.01), which is a measure of the strength of a perturbation. **Figure S3H** shows this alternative metric, which shows that the regression accurately separates strong vs. weak perturbations. Finally, **Figure S3I** compares the number of differentially expressed genes (adjusted *p* < 0.01) of the *β* coefficients computed using either all singlets in the experiment (33681) or all droplets in the experiment containing up to 6 cells, showing consistently higher numbers of genes reach significance due to the larger number of measurements enabled by droplet overloading.

### Simplified regressions for detecting on-target activation and newly expressed genes

For genes with very low expression, we cannot fit gene-level correction factors (which would be very noisy). To extend our analyses to these genes, we therefore opted for a simplified regression procedure that compensated only for the saturation effects described in the main text. We focused on a “light touch” approach to minimize distortions that might be introduced when very few events are observed.

We first constructed a modified design matrix. We begin with a binary design:

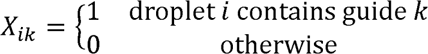

We are interested in changes relative to control cells, so we add a column that corresponds to a modified intercept that we called “control”:

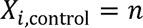

where *n* is the number of guides/cells detected in droplet *i*. Each cell within the droplet thus contributes two terms to the regression: a “control” intercept term and a perturbation driven by the guide detected. We are interested in the values of the latter coefficients, which will measure the deviations introduced by the guides to the underlying average cell state (measured by the control term).

Because we are not normalizing the data as in the previous regression approach, we had to account for the UMI saturation effect. Instead of normalizing each row to sum to 1, we computed occupancy-level-dependent scale factors. Specifically, we calculated within each gemgroup the median UMI count at each droplet occupancy level, and normalized each of these by the median UMI counts observed in singlet droplets. This produces the factors in Figure 1D. As noted in the main text, there are diminishing returns so that e.g. a droplet containing two cells does not contain twice as many UMIs, but instead ∼1.77 times as many. We use these scale factors to normalize each row of *X*. For example, if row *i* is a droplet containing 2 cells, we will scale that row by dividing by 1.77 (and not 2, which would not take the saturation into account).

With these modifications in place, we conduct the same regression as before:

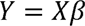

except that *Y* is now unnormalized (raw) gene expression measurements of all genes for which any counts are detected in the experiment. The non-control coefficients in *β* collect the effects induced by different guides, with *p*-values assigned as before. To prevent discoveries driven by extremely rare events, our analyses included both a *p*-value threshold and a hard “count” thresholds on the number of droplets in which non-zero counts were observed. These thresholds are described in the methods sections for the specific analyses.

### Percentile transformations, related to Figure 2E-G, Supplemental Figure 3E,I

Low UMI count regressions were used for percentile calculations and characterization of on-target guide activation. Expression was transformed into percentiles of control cell, i.e. baseline, expression range to characterize expression relative to the normal physiological range. *Preparation and normalization*: the mean expression across control cells was rounded to zero at a minimum value of 0.001 to remove noise then log10 transformed with a small constant, (10^−6^) to avoid undefined logarithmic calculations. *Construction of Reference Percentile Distribution*: To establish a reference range, we took all values greater than zero (−6, after log transformation) and created an evenly spaced distribution from min to max. This distribution was then used to arrange log-transformed expression values from 0-100. *Percentile Calculation and Interpolation*: Percentiles for control gene expressions were computed by determining the relative position of each log-transformed expression within the established linear distribution. To accommodate duplicated expression levels, we averaged percentiles associated with each unique logarithmic value. An interpolation function (`scipy.interpolate.interp1d`) was then generated from these averaged percentiles, facilitating the extrapolation of percentile values for gene expressions outside the observed range. *Analysis of on-target activation*: For each guide identity observed within the dataset, expression data for the guide target was reconstructed, rounded, and logarithmically transformed. Using the interpolation function derived from control data, we predicted percentile rankings for the target expression. The percentile results for all analyzed guides are available in **Supplemental Table 4**, which includes predicted percentiles for both control and experimental data.

### Categorization of on-target activation, related to Figure 2B-D, Supplemental Figure 4A,B

To classify the effects of guides on the target gene, several statistics were compiled for each guide including the adjusted p-value (adj_p) for the target gene, the number of cells containing target transcript (count_profile), and the percentile expression of the target gene within the control cells (ctrl_pctl) and the guide containing cells (target_pctl). The following criteria was used to classify targets as activated (is_activated) 1) *positive percentile change*: i.e. target percentile minus control percentile, indicating that the guide increased gene expression relative to the control. 2) *coefficient significance*: adjusted p-values (adj_p) were used to assess the statistical significance of changes. Only changes with adjusted p-values less than a threshold of 0.1 were classified as activated. *Guide ranking-* post-classification, guides were ranked to identify the most effective on-target activation: 1) For each gene, guides were ranked based on their target percentile value. The ranking was conducted in descending order, where the highest percentile indicated the strongest upregulation of gene expression (guide_rank). 2) The highest-ranked guide for each gene was identified (top_guide). This guide represents the most potent inducer of expression among the tested guides for that target gene, based on the analysis criteria. *Categorization Based on Expression Activation Status-* Each guide-target pair was categorized based on its activation status and baseline expression level: *1)* activated expressed: Guide-target pairs were categorized as ‘activated expressed’ if they were activated (i.e., met the above classification criteria) and the control percentile was above a threshold of 15, indicating substantial baseline expression. *2)* activated non-expressed: If a guide-gene pair was activated but the control percentile was at or below the threshold, indicating activation from a negligible baseline expression level. 3) inactive expressed: Guide-target pairs that did not meet the activation criteria but had a control percentile above the threshold were considered ‘inactive expressed,’ implying that the target is normally expressed but not significantly affected by the guide. 4) inactive non-expressed: Guide-target pairs that were neither activated nor expressed above the threshold. Guide classifications and statistics for all guides can be found in **Supplemental Table 4** and classifications are visualized in **Supplemental Figure 3E**.

### Analysis of maximum non-targeted expression, related to Supplemental Figure 4E,F

For each dataset, we isolated significant sgRNA-induced gene expression changes in transcription factors, using a predefined adjusted p-value threshold of 0.05. We then masked the guide-gene matrix at the guide-target index to exclude on-target effects, retaining only potential “non-targeted” interactions, i.e. potential downstream effects. The expression data was further filtered to ensure significant target detection by filtering for a count profile >= 3 (3 or more droplets with that guide must contain the transcript). The resulting matrix of filtered percentile expression was then processed to extract the maximum expression value for each target gene across all other guides.

### Bulk RNA-seq analysis, related to Supplementary Table 4

Data was processed using nf-core/rnaseq v3.12.0 of the nf-core collection of workflows^108^. The pipeline was executed with Nextflow v23.10.0^109^ with the following command: nextflow run nf-core/rnaseq -profile singularity --input samplesheet.csv --outdir/nextflow/results --fasta /genome_fasta/genome.fa --gtf /annotation/genes.gtf--star_index /STAR_Index_nextflow --max_cpus 24 --max_memory 256.GB. Reference files from GRCh38, release 34. Bulk transcription factor expression per cell type was compiled from salmon.merged.gene_tpm.tsv

### Epigenetic data processing and preparation of model features, related to Figure 2G, Supplemental Figure 4G,H

Raw sequencing reads for ATAC and CUT&RUN libraries were trimmed and filtered for quality (Q>15) and adapter content using version 0.4.5 of TrimGalore^110^ and running version 1.15 of cutadapt and version 0.11.5 of FastQC(https://www.bioinformatics.babraham.ac.uk/projects/fastqc/). Version 2.3.4.1 of bowtie2^111^ was employed to align reads to human assembly hg38 and alignments were deduplicated using MarkDuplicates in Picard Tools v2.16.0 (https://broadinstitute.github.io/picard/). Enriched regions were discovered using MACS2^112^ with a p-value setting of 0.001, filtered for blacklisted regions^113^. The BEDTools suite^114^ was used to create bigwig files normalized to 10 million mapped reads. The normalized bigwigs were averaged across replicates using version 3.5.4 of deepTools^115^ bigwigAverage.

Predicted TF binding bigwigs were generated using version 1.0.6 of maxATAC^65^ predict with ATAC bigwigs, TF model files downloaded from https://github.com/MiraldiLab/maxATAC_data/, and a bed file for genomic regions ±10,000 bp surrounding guide PAM positions as input.

DNA methylation data for dermal fibroblast^64^ was downloaded from NCBI GEO^116^ accession number: GSE186458 (https://www.ncbi.nlm.nih.gov/geo/query/acc.cgi?acc=GSM5652204, filename: GSM5652204_Dermal-Fibroblasts-Z00000423.hg38.bigwig). List of KRAB zinc fingers was downloaded from KRABopedia^66^ (https://tronoapps.epfl.ch/web/krabopedia/ind.php).

For each TSS, bigwig signals were averaged over a window centered at TSS (200 bp for ATAC and predicted TF tracks, 500 bp for CUT&RUN and DNA methylation). For signals other than predicted TF binding tracks, which were already scaled to between 0 and 1 by maxATAC, each was log transformed and then minmax scaled to between 0 and 1 across all TSSes.

### Categorization of TSS activation of non-expressed genes, related to Figure 2G, Supplemental Figure 4G

Genes with expression below 15 percentiles in control cells were defined as non-expressed genes. Each guide was mapped to the genome and assigned to the closest TSS. Analyses were restricted to guides mapped to up to 700 bp upstream and 300 bp downstream of a TSS. Sex chromosomes were excluded because maxATAC only makes predictions for autosomes. A TSS is defined as activatable if two out of three or at least three guides have an adjusted p-value less than 0.01 and percentile change greater than 15. A TSS is defined as non-activatable if zero guide has an adjusted p-value less than 0.01 or percentile change greater than 15 or transcriptional phenotype.

### Feature selection and evaluation of model performance, related to Figure 2G, Supplemental Figure 4L)

Logistic regression with L1 regularization was performed using version 1.4.1.post1 of scikit-learn^117^ `sklearn.linear_model.LogisticRegression` with parameters: `penalty=“l1”, solver=“liblinear”, C=0.2, class_weight=“balanced”`. Features that have non-zero coefficients were selected.

Stratified 5-Fold cross-validated predictions were made with selected features using scikit-learn `LogisticRegression (solver=“liblinear”, class_weight=“balanced”)`. For each fold, true positive rate and false positive rate were calculated using scikit-learn roc_curve. True positive rates were then interpolated at 500 evenly spaced points of false positive rates between 0 and 1. Cross-validated receiver operating characteristic (ROC) curves were created by averaging interpolated true positive rates over the folds and plotting against the evenly spaced points of false positive rates. Area under the ROC Curve (AUC) was calculated for each fold using scikit-learn `roc_auc_score` and average over the folds to give cross-validated AUCs with standard errors.

### Reduced model training on Hs27 and validation on RPE-1, related to Supplemental Figure 4N

A scikit-learn `LogisticRegression (solver=“liblinear”, class_weight=“balanced”)` model was trained on Hs27 using three features – pred-YY1, H3K27me3, and KRAB zinc fingers. The model was then used to make predictions on RPE-1 cells. The predicted probabilities were used to generate a ROC plot using scikit-learn `roc_curve` and calculate AUC using scikit-learn `roc_auc_score`.

### Comparing predictions for Hs27-specific activation, related to Supplemental Figure 4O

Genes that are uniquely activatable, by criteria described above, in Hs27 cells were selected. The reduced model above was used to make predictions in Hs27 and RPE-1 cells. The predicted activation probabilities were compared between the cell types. We could not evaluate RPE-1-specific genes because only three were uniquely activatable in RPE-1 by criteria above.

### Enrichr enrichment analysis of activable non-expressed genes, related to Supplemental Figure 4M

Enrichr^67,68^ enrichment analysis was performed on Hs27 cells using version 1.0.4 of GSEApy^118^ gseapy.enrich, with “ENCODE_Histone_Modifications_2015” as gene-set library (https://maayanlab.cloud/Enrichr/#libraries), activatable genes as input gene list, and all non-expressed genes as background gene list. Top 10 enriched gene sets were plotted using gseapy.dotplot.

### Identifying active guides by clustering, related to Figure 3, 4

We first perform the regressions as described in the section *Computing mean guide phenotypes using linear regression*. For each experiment, this produces a matrix *β* that we will orient so that rows are guides, columns are genes, and each entry denotes the effect of that guide on that gene (relative to control cells).

Due to the seed-based off-target effects discussed in the main text, we created a procedure for identifying active guides based on clustering. Our idea was that if (hierarchical) clustering was performed at a fine enough resolution, then active guides against a common target would end up clustering next to each other. This test simultaneously assesses both the strength of the phenotype (as guides must induce a phenotype that drives clustering) and its reproducibility (as two or more guides must produce the same phenotype).

To perform the clustering, we first mask the target genes within *β*: i.e., if row *i* corresponds to a guide targeting gene *j*, then we set *β_ij_* = *nan*. This is done to prevent clustering driven solely by on-target activation, which we observed for some highly inducible, highly expressed genes. We then subsetted *β* to only the columns corresponding to genes that were differentially expressed (adjusted *p* < 0.01) by at least 5 different guides. We next computed a nan-aware guide-guide correlation matrix *R* using the library NaNCorrMp, and then converted it into a distance matrix as *D*= *1* - *R*. We then clustered this matrix using hierarchical clustering (method=’average’) to produce a linkage matrix *Z* and computed inconsistency statistics (inconsistent(Z, d=5)). Our use of correlation-based comparisons (as opposed to, say, Euclidean distance) ensures that guides with varying effect sizes or incomplete penetrance can still cluster together, as in our previous work^62^.

We then clustered the linkage *Z* with fcluster at a series of different threshold values (between 0.5 and 3) of inconsistency. As the threshold gets higher, clusters merge and grow larger. At each value of the threshold, we then checked for guides that clustered based on either guide target or the seed region of the guide according to the following criteria:

- A cluster was “gene-driven” if guides targeting a common gene formed the plurality of guides in a cluster (i.e. make up at least half of the cluster)
- Similarly, a cluster was “multi-gene-driven” if guides against two or more genes formed the plurality of guides in the cluster. (This accounted for some homologous genes or functional partners that tightly clustered in effect and are labeled with slashes on Figure 4A. E.g. “GATA4/GATA6”)
- A cluster was “sequence-driven” if the majority of guides (over 75% for more than three items, or 100% for three or fewer) in the cluster had a common seed region. The common seed could be anywhere between 3 and 6 nucleotides, and we chose the longest seed that explained the cluster if more than one possibility existed.
- Finally, a small number of clusters could potentially be explained as both gene-driven and sequence-driven (e.g. if two guides against the same gene also happened to share a common seed region).

We picked the final thresholds used for clustering in a dataset-dependent manner based on inspecting the size of clusters that formed. (As the threshold was raised, clusters would merge to a point that they became incomprehensible.) For the Hs27 fibroblast dataset we used a threshold of 1.5 for routine clustering and a threshold of 1.3 for conservative analyses. For the RPE-1 dataset we used a threshold of 1.4 for routine clustering and 1.2 for conservative analyses. At these choices most clusters are very small, containing 1-4 guides.

Guides that fell into gene-driven or multi-gene-driven clusters were termed “masked_active” (i.e. they exhibited clustering despite masking of the target gene).

Finally, we also attempted to recover some perturbations for which only a single guide was active. We called these guides expanded_masked_active. They had to meet several strict criteria:

- Fell into a gene-driven or multi-gene-driven cluster as described above with a stricter threshold of 1.2
- Had on target activity detected in the analysis in *Categorization of on-target activation (related to* Figure 2*, Supplemental Table 4)*
- Did not have a bad seed sequence as described in the analysis in *Classification of Bad seeds (related to* Figure 3*, Supplemental Table 4)*

The intuition for these guides is that they cluster along guides that we know are producing distinctive phenotypes at a high level of stringency (indicated by the threshold of 1.2). In many cases they were homologous genes based on similarity of gene names: for example, *KLF4*, for which we have only a single active guide, clusters with *KLF5* in this analysis.

### Identifying seed-driven clusters, related to Figure 3

As part of our clustering analysis (*Identifying active guides by clustering (related to* Figure 3*,4)*), we conducted a parallel analysis aimed at identifying the seeds that most strongly drove clustering. To do this we computationally traversed the linkage matrix *Z* that we used for clustering. Each guide initially occupies its own cluster, and as the traversal continues these are merged into larger and larger clusters. Within each newly formed cluster of 4 or more guides we tested for the presence of a common seed sequence. We checked for all possible suffix lengths between 3 and 8 nucleotides. If a common seed was found, we recorded the minimal height of the linkage that would cause that cluster to be formed. Seeds that rapidly drove the emergence of distinct clusters thus appeared at lower threshold heights in this analysis, which we took as a measure of their strength.

### Classification of bad seeds, related to Figure 3C, Supplementary Table 4

To classify specific “bad seed” sequences the seed thresholds in RPE-1 and Hs27 cells were ranked, and the mean of the ranks taken across the cell types. The top 75 ranked sequences were taken as the “bad seeds” and corresponded to those seeds associated with sequence-based clustering across cell types. All protospacer sequences containing a bad seed at the 3’ end were classified and the specific seed match(s) recorded.

### Embedding to visualize seed-driven clustering, related to Figure S5H

We produced an embedding for all 11258 perturbations using pymde as described previously^62^. We then computed a correlation-based distance matrix, which we used to cluster the guides using HDBSCAN (`min_cluster_size=5, min_samples=1, cluster_selection_method=’leaf’`). This experiment used a dual guide sgRNA expression construct, so each cell received two sgRNAs against the same target gene. We then examined whether clusters could be explained by a common seed region (at least 70% of protospacers in the cluster terminated in a common sequence of 3 or more bases, possibly in distinct positions in the dual sgRNA expression construct). The examples we identified are annotated on the figure, though we suspect this is an undercount.

### Extracting seed-dependent effects from fitness experiments via regression, related to Figure S5

We used linear regression to look for possible seed-dependent effects in existing CRISPRa fitness screens. Our approach was to model each guide as having two possible effects: (1) on-target effects on the target gene and (2) possible off-target effects due to the seed region of the protospacer, here defined as the 5 PAM-proximal nucleotides of the protospacer. This amounts to defining two design matrices:

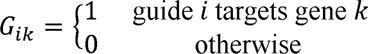

and

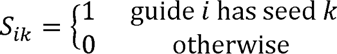

We then concatenated these two design matrices and added a constant intercept column to create the final design matrix *X*. To ensure numerical stability we used dataset-dependent filters on representation (e.g. included only guides for which 5 or more guides targeted the same target gene and included seeds as columns in *S* only if 20 or more guides had a given seed). We then regressed out experiment-dependent fitness phenotypes (different experiments used different metrics) according to:

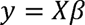

where *y* is the fitness phenotype. We assigned *p*-values using a *t*-test. Due to differences in effector, library composition, sgRNA scaffold sequence, cell type, and experimental design we do not expect the same seeds to necessarily rank the same way in all experiments. It is also unclear whether seed-dependent off-target effects necessarily always induce fitness phenotypes. Nevertheless, we observed that this procedure tended to assign certain seeds significant fitness effects, and those seeds also tended to contain PAM sequences as described in the main text. Notably the same procedure applied to a CRISPRi experiment (**Figure S5G**) did not identify strong effects. All panels with blue dots in **Figure S5** were computed according to this procedure.

For experiments that reported phenotypes for many non-targeting guides we also conducted a simpler analysis in which we grouped non-targeting guides according to seed (only for seeds that appeared in 10 or more guides), computed the median fitness phenotype, and plotted histograms (These are the yellow histograms in **Figure 3** and **Figure S5**). Non-targeting guides by construction should not have “on-target” effects, meaning any identified phenotype is likely a shared off-target effect.

The origins of the datasets used in these analyses are:

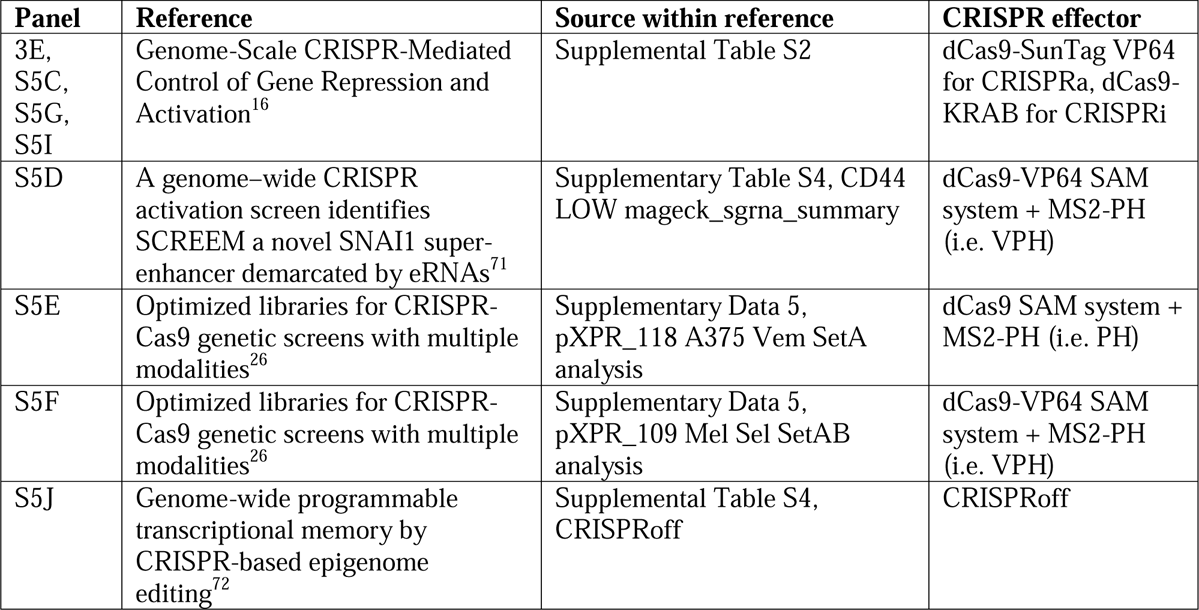

| **Panel** | **Reference** | **Source within reference** | **CRISPR effector** |
| --- | --- | --- | --- |
| 3E, S5C, S5G, S5I | Genome-Scale CRISPR-Mediated Control of Gene Repression and Activation<sup>16</sup> | Supplemental Table S2 | dCas9-SunTag VP64 for CRISPRa, dCas9-KRAB for CRISPRi |
| S5D | A genome-wide CRISPR activation screen identifies SCREEM a novel SNAI1 super-enhancer demarcated by eRNAs<sup>71</sup> | Supplementary Table S4, CD44 LOW mageck_sgrna_summary | dCas9-VP64 SAM system + MS2-PH (i.e. VPH) |
| S5E | Optimized libraries for CRISPR-Cas9 genetic screens with multiple modalities<sup>26</sup> | Supplementary Data 5, pXPR_118 A375 Vem SetA analysis | dCas9 SAM system + MS2-PH (i.e. PH) |
| S5F | Optimized libraries for CRISPR-Cas9 genetic screens with multiple modalities<sup>26</sup> | Supplementary Data 5, pXPR_109 Mel Sel SetAB analysis | dCas9-VP64 SAM system + MS2-PH (i.e. VPH) |
| S5J | Genome-wide programmable transcriptional memory by CRISPR-based epigenome editing<sup>72</sup> | Supplemental Table S4, CRISPRoff | CRISPRoff |

### Embedding of active guides, related to Figure 4A

We produced an embedding of all masked_active or expanded_masked_active guides using pymde as described in our previous work^62^. The labels are placed for (1) all multi-gene-driven clusters (which have gene names separated by slashes) (2) the top 100 genes (with guides identified as masked_active or expanded_masked_active) according to number of differentially expressed genes (adjusted *p*-values < 0.05).

### Examining conservation of perturbation effects across cell context, related to Figure 4B

We first identified perturbations that induced phenotypes in both cell types. We only considered guides that were either masked_active or expanded_masked_active in both cell types and that did not have bad seeds. For each gene we then picked a single representative guide as the “primary guide” as the one that had the highest mean (log) number of differentially expressed genes across the two cell types. This led to 121 guides in total that had phenotypes in both cell types.

We then extracted the expression profiles for these guides from the *β* matrices for each cell type produced by our regressions. We only considered genes (columns of *β*) that were chosen as part of components by our sparse PCA gene program extraction procedure in both cell types. This led to a total of 1702 genes. As in the clustering analysis, we masked the targets of guides using *nan* to prevent effects driven by CRISPRa on-target activity.

We then compared guides in two ways. “Profile correlation” corresponded to the Pearson correlation between the expression profile in Hs27 cells vs. that in RPE-1 cells. Next, we computed guide-guide correlation matrices in each cell type. We treated the rows of these matrices as “correlation profiles” for each guide. The “cophenetic correlation” corresponded to the Pearson correlation between the correlation profile in Hs27 cells vs. that in RPE-1 cells. These two perspectives essentially compare whether a guide’s absolute effects on gene expression (profile correlation) are preserved across cell context vs. whether a guide’s relationships with other guides are preserved (cophenetic correlation).

### Identifying transcription factors that oppose each other, related to Figure 4C

We reasoned that our gene program profiles (Figure 4A, bottom right) represented more meaningful representations of perturbation function. We took the gene program profiles for guides that were either masked_active or expanded_masked_active and collapsed them to gene level by averaging according to target gene. We then computed the gene-gene correlation matrix. We reasoned that strong negative associations in this matrix derived from transcription factors that might functionally oppose each other. (Strong positive associations are more difficult to reason about, as they arose frequently from homologous transcription factors.) We viewed an association as particularly strong if it fell in the bottom 0.1% of entries in the correlation matrix. We constructed a graph using these links. Figure 3C simply plots the 3 connected components in this graph that contained 3 or more transcription factors. Arcs in the circos plot annotate strong negative or positive associations (in the top 0.1% of entries of the correlation matrix) among the selected transcription factors.

### Extracting gene expression programs using consensus sparse PCA, related to Figure 4

We developed a procedure for extracting gene expression programs using sparse PCA. Interpretability is aided by a modification we made to sklearn’s `SparsePCA` class to enforce positivity of the coefficients that define the programs. (This is a built-in functionality of sklearn’s dict_learning method, which `SparsePCA` uses to extract components. We simply set `positive_code=True`.) This modification forces the modified class, which we called NonNegativeSparsePCA, to pick components that are composed of genes that move up or down together. We thus retain the interpretability benefits of methods like NMF, while adding a sparsity constraint that makes programs smaller.

We apply this method to the (unmasked) *β* matrix from the *Identifying active guides by clustering (related to* Figure 4) section. We subsetted *β* to rows corresponding to masked_active guides identified by clustering, and to columns that were differentially expressed (adjusted *p* < 0.01) by at least 2 different masked_active guides. To identify the correct number of programs to extract, we used a consensus procedure. We produced 100 bootstrap resamples of our subsetted *β* matrix and fit NonNegativeSparsePCA models to each of them. In each case we extracted 100 components and, based on testing, set the sparsity parameter alpha=1. As a result we extracted 10,000 components in total across all resamples.

To identify components that appeared recurrently across resamples, we then borrowed a consensus building procedure from Leland McInnes’s enstop repository (https://github.com/lmcinnes/enstop/). We embedded the 10,000 extracted components into a common embedding using UMAP (n_neighbors=15, n_components=5, metric=hellinger). We then cluster these embedded components using HDBSCAN (min_samples and min_cluster_size were both set at 80, meaning the component must appear in 80% or more of the 100 bootstrap resamples). In Hs27 fibroblasts, this produced 57 clusters. We then finally reran NonNegativeSparsePCA on the original subsetted *β* (i.e., containing all masked_active guides) with n_components set to 57 to ensure deterministic results. The resulting components varied drastically in size, with some containing hundreds of positive coefficients and others concentrating on only a handful of highly variable genes, showing the versatility of this procedure to extract programs of different scales.

Finally, to produce representations of guides’ effects as linear superpositions of programs, we used the transform method of NonNegativeSparsePCA (inherited from SparsePCA), which fits program scores for each guide using ridge regression. We fit these scores for all guides (but used only masked_active guides to derive the programs).

### Enrichment analysis associating fibroblast gene programs with in vivo fibroblast states, related to Supplemental Figure 10A

We performed permutation testing to assess the degree of overlap between genes in the CRISPRa programs and in vivo fibroblast states defined by Korsunsky *et al.*, 2022^54^. The genes for each gene program were defined as the top 50 genes for each program based on sparse PCA. The genes for each cluster were defined as any differentially expressed gene in that cluster keeping each tissue separate. The Jaccard similarity was computed between the program genes and cluster genes. Permutations were generated by concatenating the program genes together and randomly sampling sets of program genes equivalent in size to the program. This method allows for duplicate genes which are removed when computing Jaccard similarity. A total of 1,000 permutations were performed for each program-cluster combination. P-values were computed by computing the number of permutations with a Jaccard similarity greater than or equal to the observed Jaccard similarity and dividing by the total number of permutations. Overlap coefficients were computed by taking the number of shared genes between the cluster and program and dividing by the number of program genes. Permutation-based p-values were then adjusted using the Benjamini-Hochberg correction for multiple hypothesis testing. A FDR threshold of 0.1 was used to determine significant cluster-program links.

### Analysis of newly expressed genes, related to Figure 5D-E, Supplemental Figure 10D

Newly expressed genes for each guide were defined from Low UMI count regressions as those showing a positive percentile change (pctl_change), with an adjusted p-value less than 0.01, the number of cells containing target transcript (count_profile) greater than or equal to 3, and the percentile expression of the target gene within the control cells (ctrl_pctl) as less than 15. Counts of guides per target activating each gene were used to filter for high confidence new expression.

### Curation of cell state markers, related to Figure 5C, E-F, Supplemental Figure 10C-D

*Fibroblast cell states*- Supplementary data tables for human fibroblast clusters were downloaded from Buechler and Pradhan *et al.,* 2021^53^ (Supplementary Table 7, “All markers”) and Korsunsky, 2022^54^ (Supplemental information, Document S2, Supplementary Data,“TableS8”). From the Buechler and Pradhan *et al.,* 2021 dataset, the annotated cluster 8 (human universal state/normal) and cluster 3 (inflammatory/cancer-associated fibroblasts) were isolated and filtered for avg_logFC > 1 and p_val_adj < 0.05. For marker genes from Korsunsky *et al*., 2022 clusters “CD34+MFAP5+ C9” and “SPARC+COL3A1+ C4” were isolated. Clusters were further subset by Tissue classification and two gene lists generated one for all Tissue classes except “Marginal” and the other limited to the Marginal class (i.e. cluster specific effects regardless of tissue of origin). As described within Korsunsky *et al*., 2022, markers that were significant across all 4 tissues—SalivaryGland, Lung, Synovium, and Gut—were filtered as high confidence cluster markers. The set of significant genes for the Marginal class were also incorporated as part of our marker gene sets. Pval < 0.05 and LogFoldChange > 1 was used to filter significant marker genes and the top 100 genes from each cluster selected. The final marker gene list was further supplemented with selected of differentially expressed fibroblast markers from Smith *et al*., 2023^40^ Fig. 1b.

*Retinal markers*- known retina marker genes were compiled from the Human Protein Atlas^90^ retina proteome (https://www.proteinatlas.org/humanproteome/tissue/retina) and computational selected from the RNA FANTOM tissue gene data (https://www.proteinatlas.org/download/rna_tissue_fantom.tsv.zip). Selection was performed by first calculating a specificity score for each gene within all tissues. First, all rows and columns of the expression matrix were normalized to have the same sum though an iterative procedure. Then, a specificity score was assigned to each gene within each tissue as the ratio of expression within the tissue over the sum of all tissues. The top 500 retina specific genes (highest ratio) were selected as retinal markers. Additional cell type specific markers were hand compiled from the Single Cell Type Atlas^2^ (https://www.proteinatlas.org/humanproteome/single+cell+type) curated sets of cell type markers including glial types (Muller glia cells, Schwann cells) and pigmented cells (Melanocytes).

### Visualization of fibroblast marker activation and variation, related to Figure 6F, S10C,D

Initially, lists of high confidence gene activation (> 2 guides per target) for outlier perturbations (i.e. with many genes activated) were compared to known marker genes for fibroblast state including Buechler and Pradhan *et al.*, 2021: universal clusters - PI16, DPT, COL15A1, defined steady state clusters - COMP, CCL19, COCH, CXCL12, FBLN1, BMP4, NPNT, HHIP, Korsunsky *et al*. 2022 named clusters: CD34/MFAP5, CXCL10/CCL19, FBLN1, PTGS2/ SEMA4A, SPARC/COL3A1. Subsequently, high confidence gene activation for all targets was cross referenced with the full curated list of cell state markers. This process was similarly applied to newly expressed genes from RPE-1 perturbations, cross referencing with retinal marker genes and curated cell type markers.

### Associating transcription factors with markers of *in vivo* phenotypes, related to Figure 6A,B

In this analysis we were took a more inclusive view of which perturbations to include than when extracting gene programs. We considered all guides that: (1) were not in seed-driven clusters in either Hs27 or RPE-1 datasets; (2) did not contain bad seeds that we had identified; and (3) were either masked_active, expanded_masked_active, or showed detectable on-target activation in either Hs27 or RPE-1 cells. 5346 guides passed these filters. To collapse these to gene-level program scores, we for each program averaged the two guides with the strongest absolute effects for that program. (A large negative score by one guide could therefore potentially cancel a large positive score by another.) This led to predictions for 1665 transcription factors. For the universal state we used the score of program 31 as the *in vitro* score and for the inflammatory state we used the score of program 32. These scores form the *y*-axis of **Figure 6A,B**.

We then compared these scores with *in vivo* expression of state markers for these two states. We downloaded rna_single_cell_type_tissue.tsv.zip from the Human Protein Atlas website, which aggregates single-cell RNA sequencing data about 81 cell types in 31 distinct datasets into clusters of single cells that have been averaged together. Each cluster is annotated with a cell type. We included clusters annotated as either “fibroblasts”, “smooth muscle cells”, or “endometrial stromal cells.” We removed clusters derived from the “Early spermatids”, “Leydig cells”, and “Late spermatids” tissues as they exhibited markedly distinct expression characteristics from other datasets. We then normalized the expression profiles of all clusters to sum to 1, and *z*-normalized each gene feature. The resulting dataset contains 96 cluster gene expression profiles across 20082 genes.

We extracted the expression of specific state markers from this dataset: *PI16* for universal fibroblasts and *SPARC* for inflammatory fibroblasts. To measure similarity of expression patterns, we then correlated marker expression with the expression of each of the 1665 transcription factors within the dataset. These scores form the *x*-axis of **Figure 6A,B**.

Outliers were identified independently for each variable using the 1.5 interquartile range (IQR) rule, where the IQR is the difference between the first quartile (Q1, 25th percentile) and the third quartile (Q3, 75th percentile). Data points below Q1 - 1.5 * IQR or above Q3 + 1.5 * IQR were considered outliers. Positive outliers were defined as data points above Q3 + 1.5 * IQR. We highlighted data points that were positive outliers in both variables simultaneously, as they were both (1) capable of inducing the state (*y* outlier) and (2) strongly correlated with the state marker *in vivo* (*x* outlier).

### Scoring of transcription factor phenotypes for similarity to *in vivo* observed transcriptional states, related to Figure 6C

We used the consensus fibroblast cell state lists from from Korsunsky *et al*., 2022^54^ (see *Curation of cell state markers*) to select genes for scoring, intersecting with the expressed genes used within linear regression of mean guide phenotypes (columns of *β*). Guides were then filtered for those within the “final library” (i.e. not seed driven, and observed as having on-target activity within either cell type). The top 2 guides for each target were selected based on the highest number of differentially expressed genes observed for each guide (“de_genes_fibro_final”). The mean expression profiles for each target were then generated by taking the average of the *β* coefficients across the selected guides, and on-target effects were masked. Extreme gene variation (i.e. z-score > 4) was capped at a maximal variation of 4 to limit the score contribution from any one outlier gene. For each, “Cluster” and “Tissue” within the differentially expressed gene list from Korsunsky *et al*., 2022^54^ (Supplemental information, Document S2, Supplementary Data,“TableS8”) we used the differentially expressed genes within that “Tissue” to create a weighted score of transcription factor phenotype. Specifically, we multiplied each gene’s expression deviation by the “LogFoldChange” observed for each “Tissue” and a final score was created by summing the weighted deviations dividing by the total weight (sum of the absolute “LogFoldChange” per “Tissue”). In the case of the MYH11+ C13: Synovium very few differentially expressed genes (<10) overlapped with the consensus gene list for the MYH11+ C13 cluster (consistent with the previously observed absence of the myofibroblasts from the synovium^54^) and the tissue was dropped from the analysis. To visualize the top scoring transcription factors per “Cluster”, each “Tissue” nominated it’s top 2 scoring transcription factors to be visualized regardless of transcription factor expression within the “Tissue”. Mean expression (mean log normalized counts per cell, generated from ported Seurat object) from each cluster for each nominated transcription factor was used to annotate the visualization, with the highest expressed transcription factor marked in **bold** and those with observed expression > 50^th^ percentile of all transcription factor expression marked with *italic*.

### Processing of the Ovarian cancer fibroblasts, related to Figure 6D

Annotated fibroblasts from the Vázquez-García *et al.,* 2022 MSK-SPECTRUM dataset were processed from raw counts. Briefly, the fibroblast subset was filtered in scanpy with `scanpy.pp.filter_genes(adata, min_cells=3)`, the dataset was previously filtered for minimum genes >500, >1,000 UMI counts, and <25% mitochondrial genes^104^. Counts were normalized to median total counts and Log(x+1) scaled with `scanpy.pp.normalize_total(adata)` and `scanpy.pp.log1p(adata)` respectively. Highly variable genes were selected with sc.pp.highly_variable_genes(adata, n_top_genes=2000, batch_key=“batch”). Batch correction was performed with bbknn^119^ using the workflow from Park *et al.*, 2020^120^. Specifically:

~~~
bbknn.bbknn(adata)
scanpy.tl.leiden(adata, resolution=0.4)
bbknn.ridge_regression(adata, batch_key=[’batch’],confounder_key=[’leiden’])
scanpy.tl.pca(adata)
bbknn.bbknn(adata)
~~~

We observed that patient sample “SPECTRUM-OV-045” exhibited poor batch correction evidenced by separation upon visualization of data integration, `scanpy.pl.umap(adata, color = [“batch”]),` and this sample was removed from further analysis. *PI16* and *KLF4* expression was visualized with `scanpy.pl.umap(adata, color=[’PI16’, ‘KLF4’])`.

### CIRCOS plot of program cross regulation, related to Figure 6E

Gene-level program scores were calculated as described in *Associating transcription factors with markers of in vivo phenotypes,* allowing for program scores from all transcription factors with either observed phenotypes or on-target activity. The top 10 program activating (i.e. highest program scores, positive) and repressing (lowest program scores, negative) were selected for programs 4 (myofibroblast), 31 (universal), and 32 (inflammatory) and visualized with `pycirclize.Circos` (https://github.com/moshi4/pyCirclize) with `track.heatmap` for each TF’s program score and links (`circos.link`) drawn between transcription factors recurring across the programs. Links were visualized as “red” if both program scores were positive, “blue” if both were negative, and “purple” if the values were opposing (i.e. program cross-regulation).

### Selection of Final Filtered guide library, related to Supplementary Table 4

The final validated guide library was selected from the sets of “active” and “expanded_active” classified guides for each cell line, as well as guides that showed direct on target activation (i.e. “is_activated”). All guides were filtered for seed driven clustering (“seed_driven”) or the presence of a categorized “bad_seed” within the protospacer sequence. The final guide library can be found in **Supplemental Table 4**.

