## Supplementary material for "Comprehensive transcription factor perturbations recapitulate fibroblast transcriptional states": Document S1 Figures S1-S9 and Supplemental Figure legends

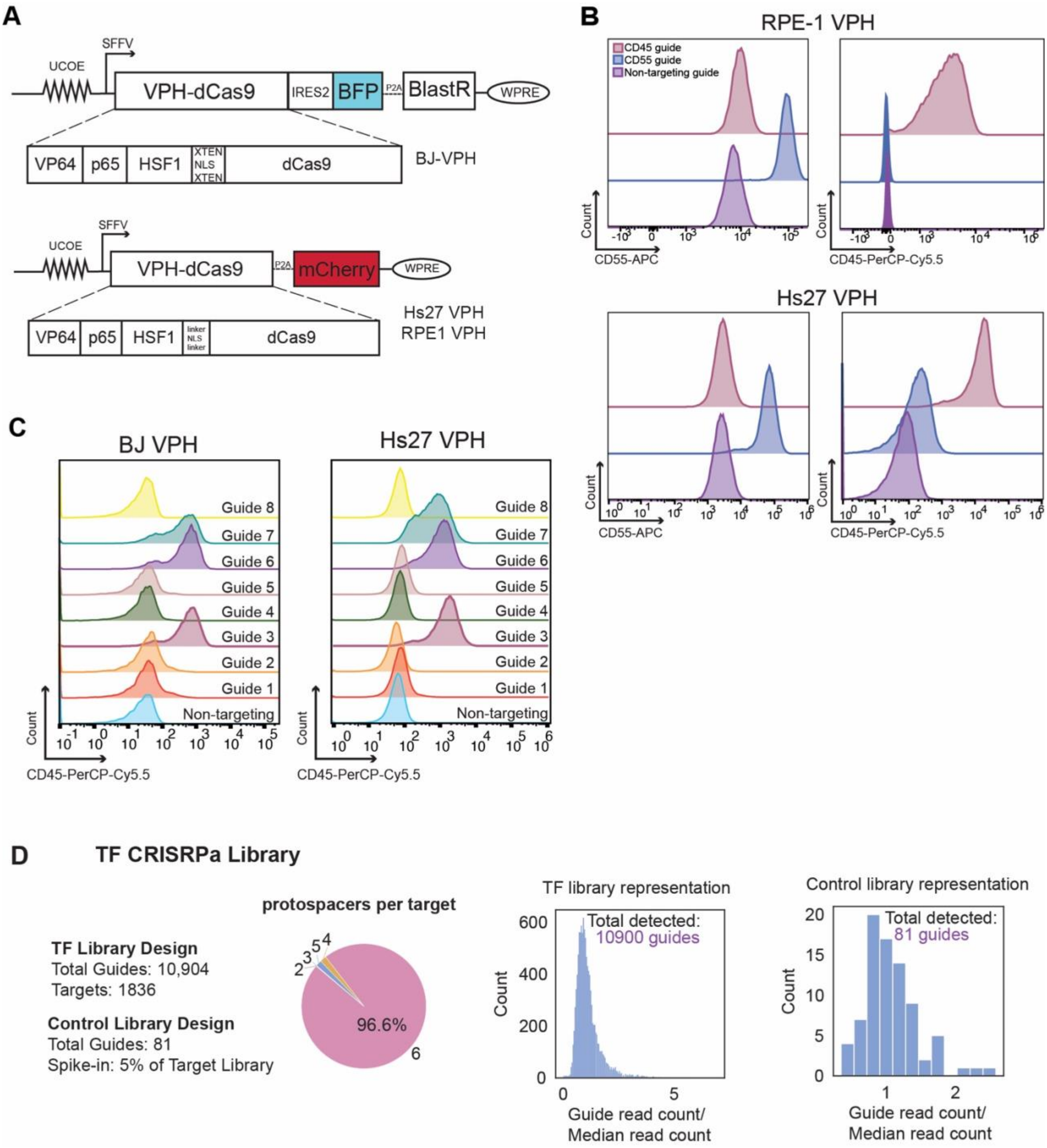


**Figure S1. Development and testing of VPH-dCas9 CRISPRa effector.**

A. Schematic of VPH-dCas9 CRISPRa effector constructs used in this study. BFP construct (top) was used in BJ creating BJ fibroblast CRISPRa cell lines, while the mCherry containing construct (bottom) was used in to create Hs27 fibroblast and RPE-1 CRISRPa cell lines.
B. Flow cytometry data demonstrating activation of cell surface markers CD45 and CD55 by the VPH-dCas9 CRISPRa system in Hs27 and RPE-1 cells.
C. Flow cytometry data showing variable efficacy of predicted CRISPRa guides targeting *CD45* in BJ and Hs27 fibroblast cells.
D. Description of transcription factor-CRISPRa (TF-CRISPRa) library design and quality control.

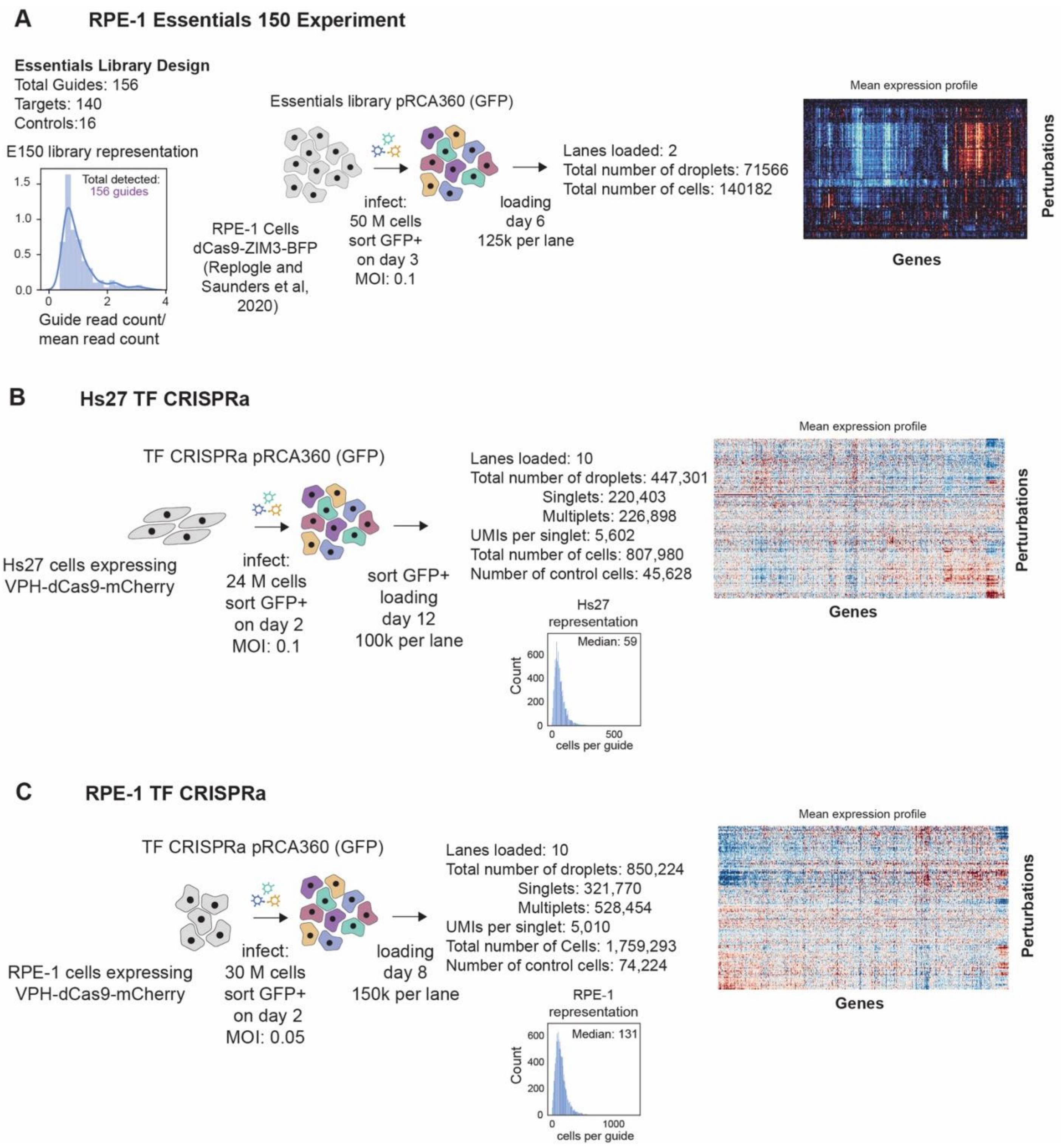


**Figure S2. Experimental design parameters for the major experiments in the paper.**

A. Schematic and features of benchmarking experiment testing knockdown of essential genes in RPE-1 cells using CRISPRi.
B. Schematic and features of CRISPRa experiment targeting all transcription factors in Hs27 fibroblast cells. Bottom: distribution of guide frequencies in Hs27 (i.e. the number of cells per guide). Due to our use of droplet overloading, some cells containing a guide will end up co-encapsulated in droplets with other cells.
C. Schematic and features of CRISPRa experiment targeting all transcription factors in RPE-1 retinal pigment epithelial cells. Bottom: distribution of guide frequencies in RPE-1 (i.e. the number of cells per guide). Due to our use of droplet overloading, some cells containing a guide will end up co-encapsulated in droplets with other cells.

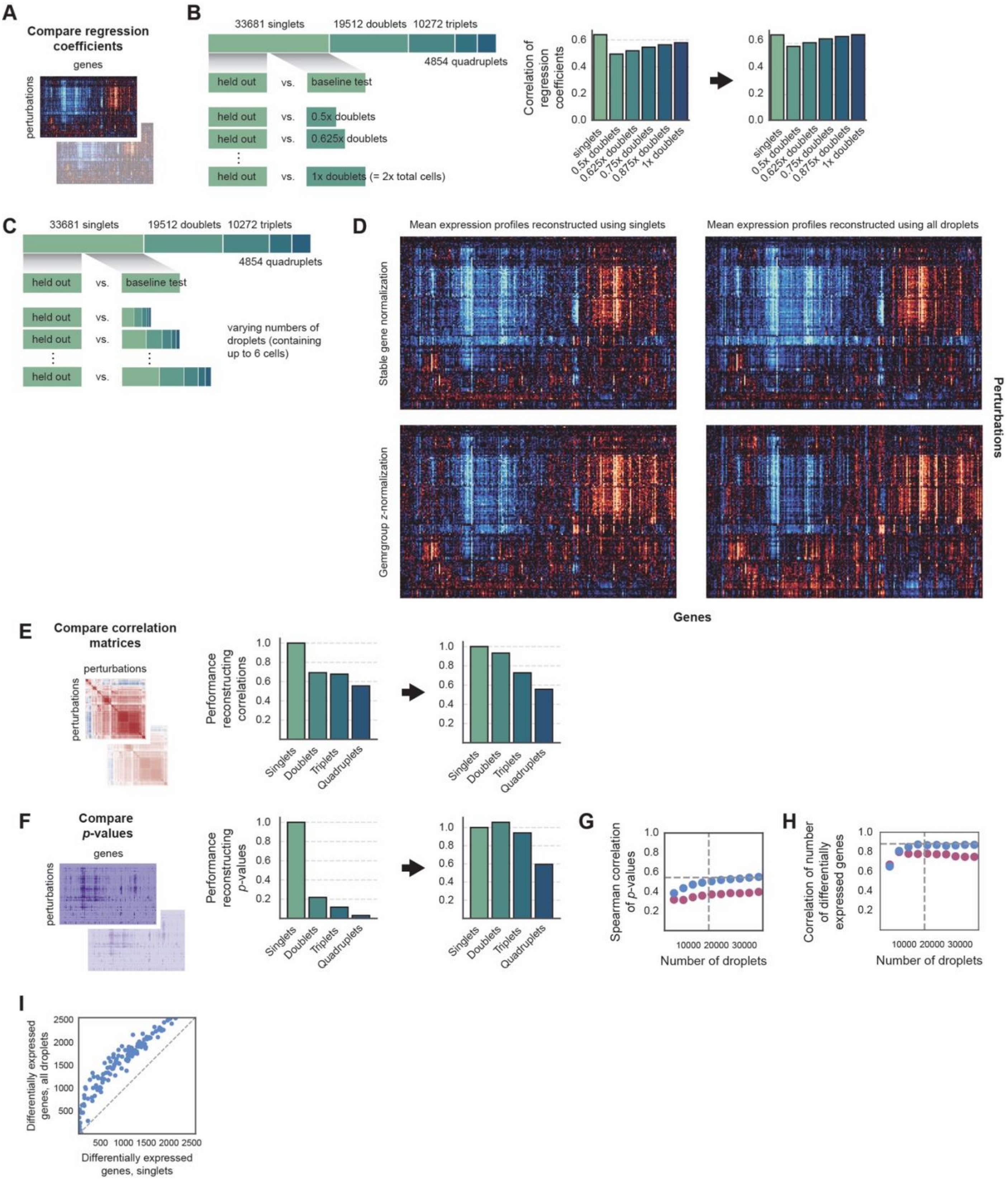


**Figure S3. Testing performance of droplet overloading model, related to Figure 1**

A. In computational benchmarking experiments, we separate the dataset into disjoint subsets of various types, infer regression coefficients measuring effect sizes using our linear regression model, and then compare reconstruction accuracy using several different metrics. The simplest metric compares the overall structure of coefficients by computing the correlation coefficient between the (stacked) regression coefficient matrices.
B. Design of doublet downsampling experiment. We first establish a baseline by comparing the correlation of regression coefficients between two disjoint sets of 16,800 singlets. To assess performance reconstructing coefficients using only doublets, we then subsample the 19,512 doublets in the experiment to varying depths and compute the correlation of regression coefficients (right).

C. An alternative experiment design reflects how the approach is used in practice, by subsampling mixtures of droplets proportionally to how often they appear.
D. Visual comparison of regression coefficients inferred using all withheld singlets (left) vs. all withheld droplets containing up to 6 cells. Results are compared with our stable gene-based normalization (top) vs. without (bottom). Stable gene normalization leads to visually indistinguishable results, while baseline normalization causes structural differences to appear driven by differences in transcript capture efficiency.
E. Another metric for comparing reconstruction accuracy is to instead compare the perturbation-perturbation correlation matrices, which summarize relative comparisons between perturbations that tend to drive functional analyses such as clustering. This experiment follows the same design as in Figure 1H.
F. Finally, the regression procedure assigns *p*-values to differentially expressed genes. We compare the structure of these matrices by stacking them and computing Spearman correlations. This experiment follows the same design as in Figure 1H.
G. Spearman correlation of *p*-values using the experiment design from panel (C). Blue and magenta dots denote performance with and without stable gene normalization, respectively. Dashed lines show performance using disjoint set of 16,800 singlets.
H. An alternative metric is to summarize each perturbation according to the number of differentially expressed genes (at a false discovery rate of 5%) and compute the Spearman correlation between different reconstructions. This test assesses whether reconstructions order the overall strength of perturbations correctly in different sampling regimes. Blue and magenta dots denote performance with and without stable gene normalization, respectively. Dashed lines show performance using disjoint set of 16,800 singlets.
I. Comparison of the number of the number of differentially expressed genes (at a false discovery rate of 5%) discovered using only singlets vs. all droplets containing up to 6 cells. Dashed line: line of equality.

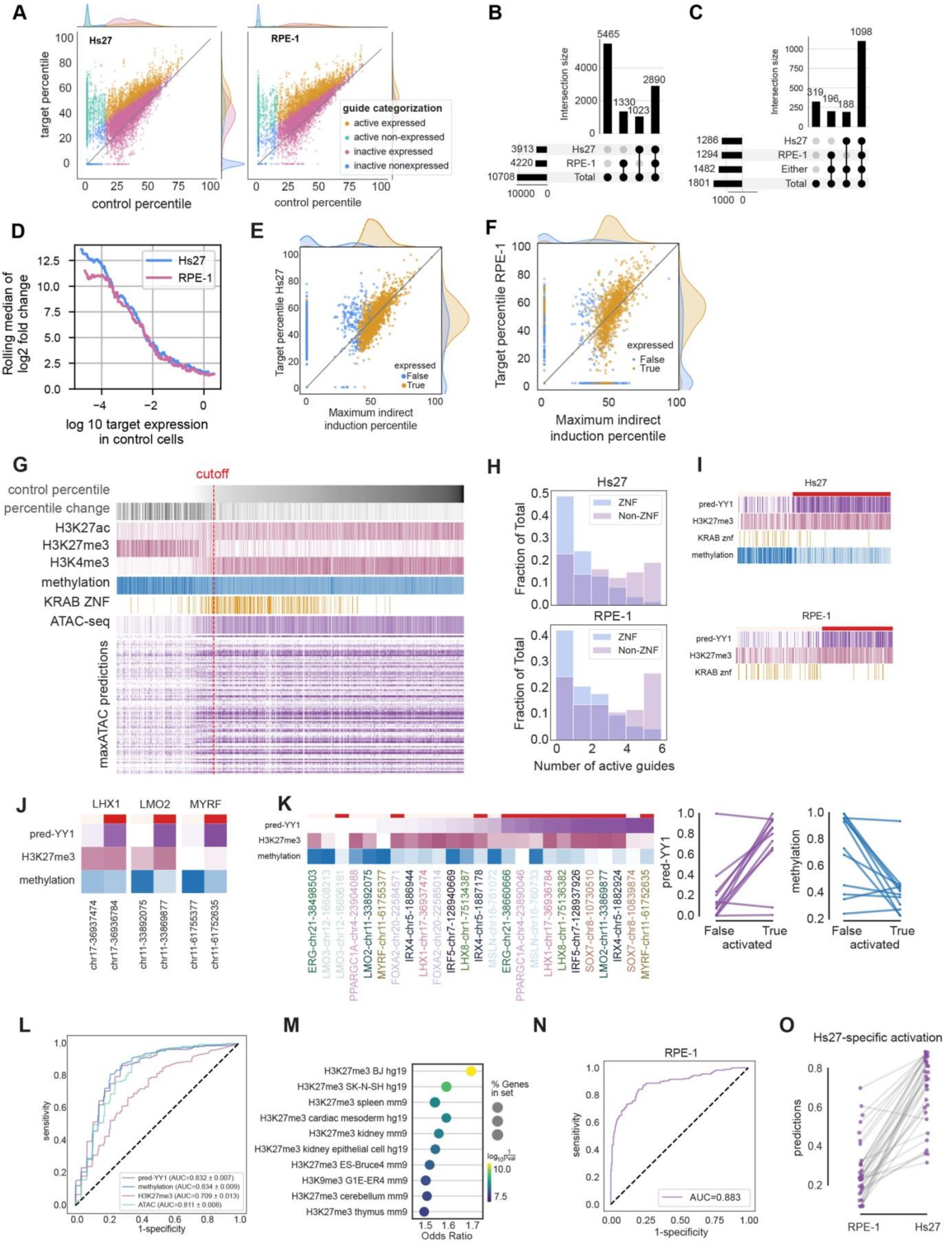


**Figure S4. On-target activity of the TF-CRISPRa library and modeling of the epigenetic determinants of susceptibility to CRISPRa, related to Figure 2**

A. Detailed summary of target gene activation in Hs27 vs. RPE-1 cells, for comparison with Figure 2C. Here we categorize guide efficacy according to whether the target gene is expressed in control cells (containing non-targeting guides) and whether the guide is determined to be active according to our regression procedure

(Methods). The procedure is conservative given the noise intrinsic to single-cell RNA sequencing so some guides exhibiting marginal activity will be called as inactive.

B. Upset plot of guide activity across Hs27 and RPE-1 cells.
C. Upset plot of activatable target genes across Hs27 and RPE-1.
D. Diminishing returns of CRISPRa activation as target gene expression level increases. Genes were ordered by mean expression level in control cells and a rolling median of the $\log_2$ fold activation was computed.
E. Comparison of CRISPRa-induced activation of the on-target transcription factor to the maximal observed activation the experiment as an indirect consequence of activating another transcription factor (Hs27).
F. Comparison of CRISPRa-induced activation of the on-target transcription factor to the maximal observed activation the experiment as an indirect consequence of activating another transcription factor (Hs27).
G. Selection of target promoters used for examining impact of epigenetic factors on amenability to CRISPRa. We focused our attention on target genes that were not expressed in control cells containing non-targeting guides and that were activatable by at least two out of three available guides or at least 3 distinct guides per target. Plot compares the epigenetic features used for model training for genes above and below our expression threshold.
H. Recalcitrance of KRAB zinc finger proteins to activation by CRISPRa. Plot compares number of active guides for KRAB zinc finger transcription factors vs. other transcription factors.
I. Top: Selected features at promoters that were and were not susceptible to activation in Hs27. Bottom: Selected features of reduced model at promoters in RPE-1 cells.
J. For a subset of target genes, our experiment included guides targeting distinct transcription start sites that behaved differently. Plots compare selected features at example promoters in Hs27 cells. Red indicates that the promoter is activatable.
K. Comparison of model features at alternative promoters of all genes where CRISPRa selectively activated only one of the two promoters (see also J). Red indicates that the promoter is activatable. Right: Summarization of pred-YY1 and methylation signals at pairs of promoters according to activatability.
L. Performance of logistic regression models trained using individual epigenetic features to predict susceptibility to CRISPRa in Hs27 cells.
M. Enrichr enrichment analysis of genes susceptible to CRISPRa in Hs27 cells using the "ENCODE_Histone_Modifications_2015" gene set. The top hit is H3K27me3 in BJ cells, a primary fibroblast cell line similar to our Hs27 cells.
N. Predictive performance in RPE-1 cells of a reduced model trained on pred-YY1, H3K27me3, and KRAB zinc finger features from Hs27 cells.
O. Comparison of model scores in RPE-1 and Hs27 for genes only activatable in Hs27 cells.

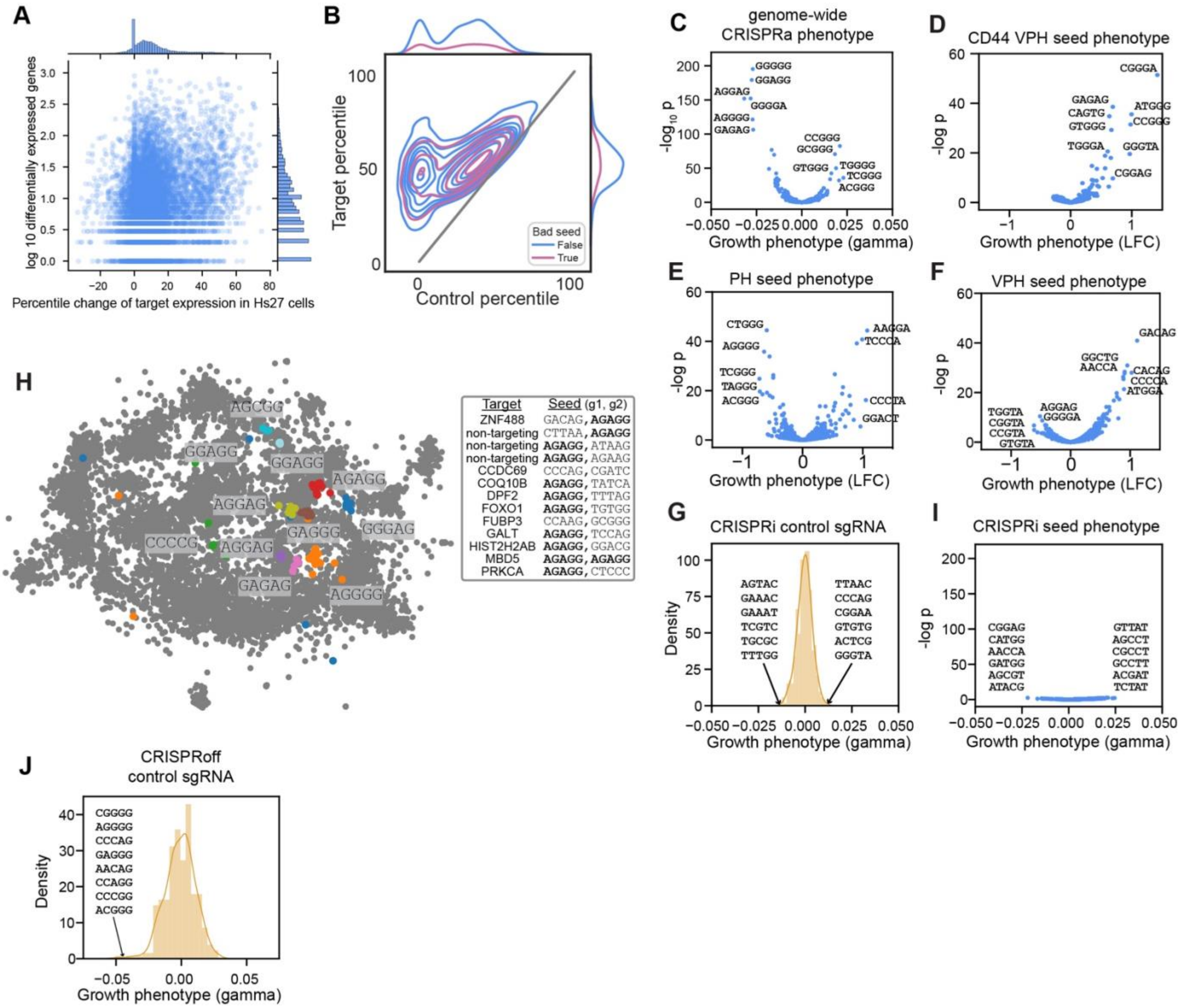


**Figure S5. Seed-driven phenotypes in other datasets, related to Figure 3.**

A. Scatter plot comparing the strength of on-target activation induced by each guide to the number of differentially expressed genes.
B. Comparison of target activation between guides with or without bad seeds.
C. In a genome-wide CRISPRa fitness screen using an alternative SunTag CRISPRa effector[16], a regression model (Methods) assigning guides' effects to on-target and seed-dependent components identifies seeds with consistent fitness effects across guides.
D. Regression analysis as in C applied to a CRISPRa screen targeting enhancers using an alternative VPH CRISPRa effector[71].
E. Regression analysis as in C applied to a genome-wide CRISPRa using a PH CRISPRa effector[26].
F. Regression analysis as in C applied to a genome-wide CRISPRa using an alternative VPH CRISPRa effector[26].
G. Median fitness effects in non-targeting control sgRNAs grouped by seed in a genome-wide CRISPRi screen[16].
H. Identification of seed sequence-driven clusters in a large-scale CRISPRi Perturb-seq experiment in K562[62]. In this case each construct contains two guides (guide seeds shown as: g1, g2), with one guide almost always containing an AGAGG seed.
I. Regression analysis as in C applied to sgRNAs from a genome-wide CRISPRi screen[16].
J. Median fitness effects in non-targeting control sgRNAs grouped by seed in a CRISPRoff fitness screen[72].

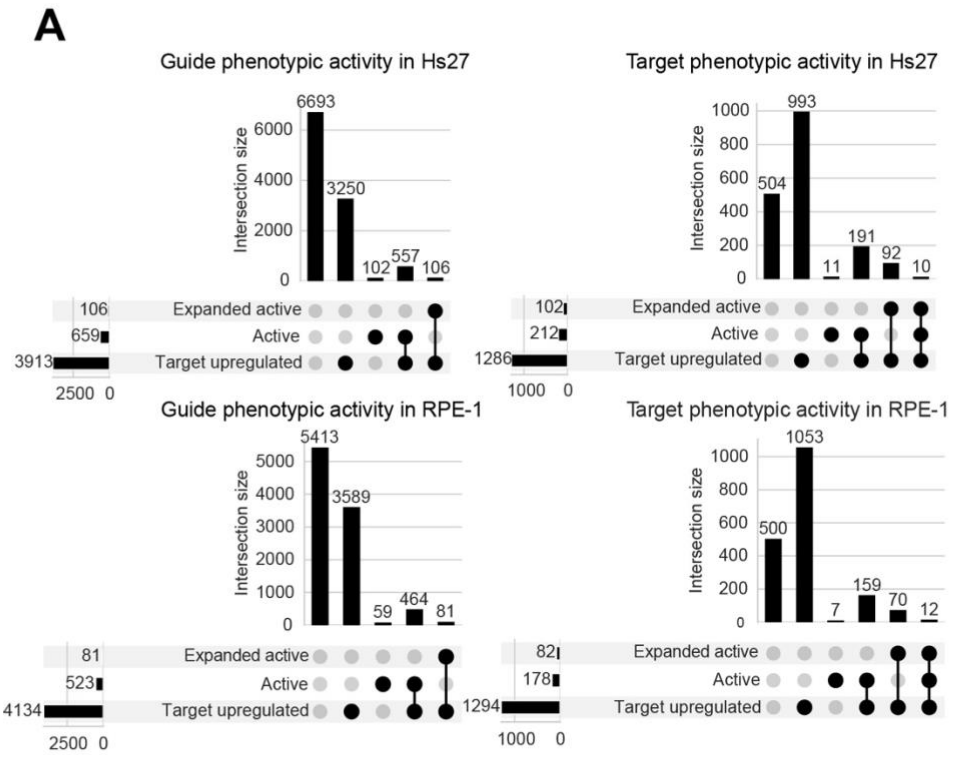


**Figure S6. Classification of guides and targets by phenotypic effect, related to Figure 4A**

A. We identified transcription factors causing reproducible changes in transcriptional phenotypes using the clustering-based criterion described in the text: at least two guides targeting the same gene must cluster together. These we term "active" guides. Because the clustering leading to this classification is fine-grained, we additionally include any guides for other transcription factors that fall into clusters marked active. These we term "expanded active" guides. Plot compares these properties of guides to on-target activation statistics from Figure 2. On-target activation for some guides will fail to be detected because our procedure for doing so is conservative.

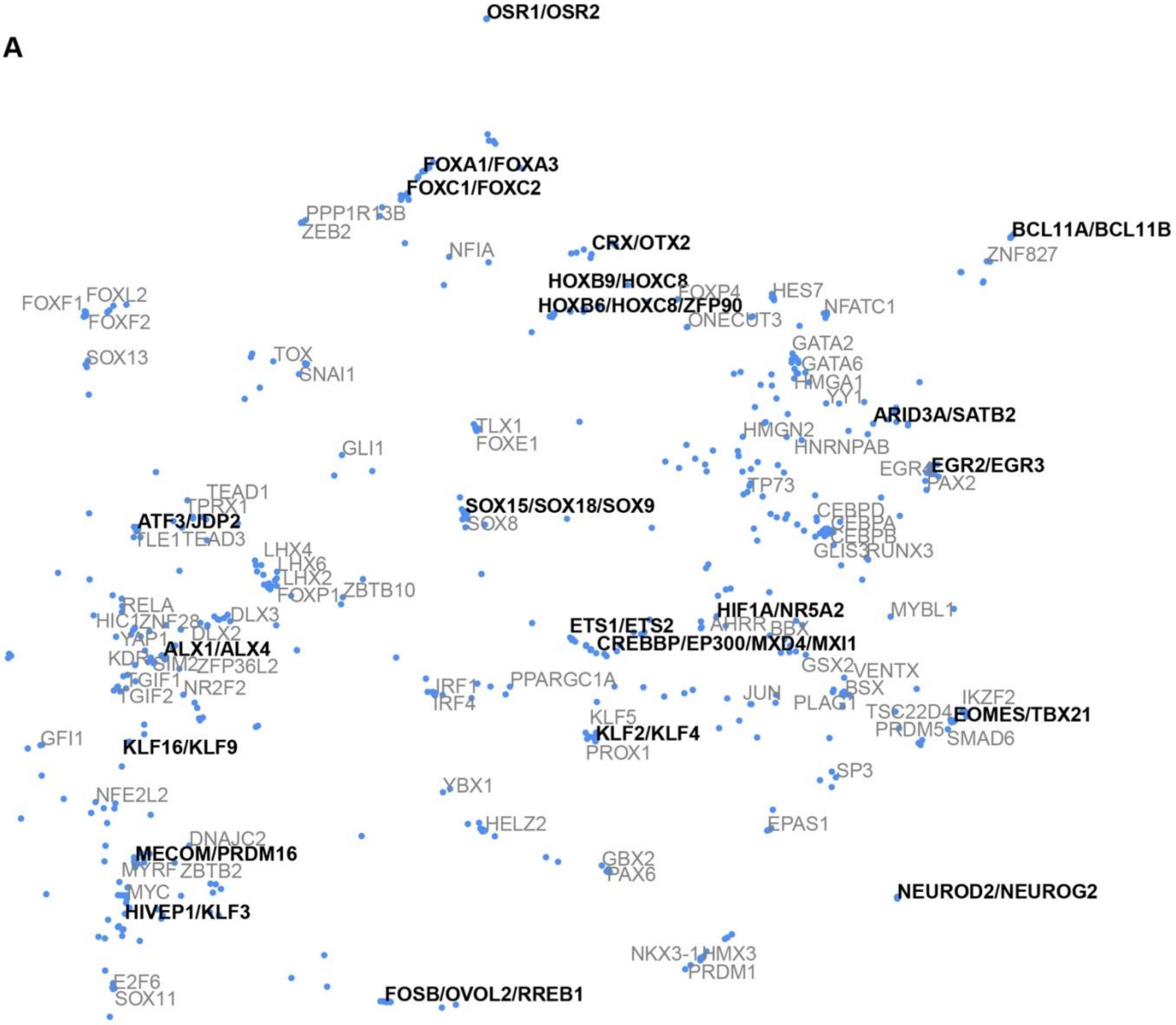


**Figure S7. Embedding of active RPE-1 perturbations, related to Figure 4A**

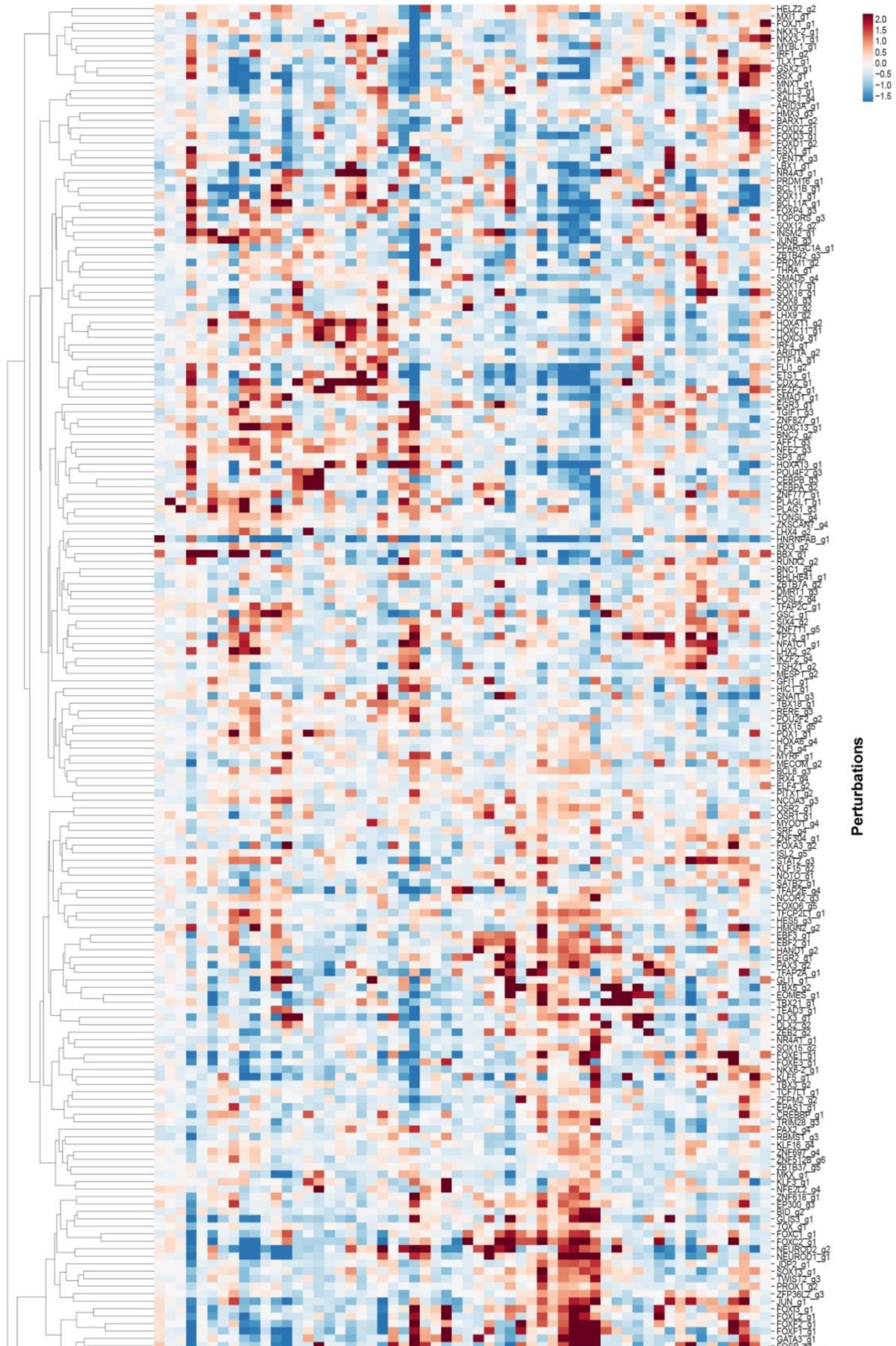

2.0
1.5
1.0
0.5
0.0
−0.5
−1.0
−1.5
Perturbations
6 36 57 12 13 29 15 56 32 53 3 30 2 23 42 11 16 24 44 17 9 38 35 47 1 28 10 21 14 22 41 27 25 4 54 51 49 46 50 52 5 0 20 33 39 45 37 7 34 26 8 43 18 19 40 48 31 55
Programs

**Figure S8. Encodings of Hs27 perturbations in terms of sparse expression programs, related to Figure 4**

A. Figure shows heatmap of inferred program activation levels for representative active guides for all target genes that were active according to our clustering-based test.

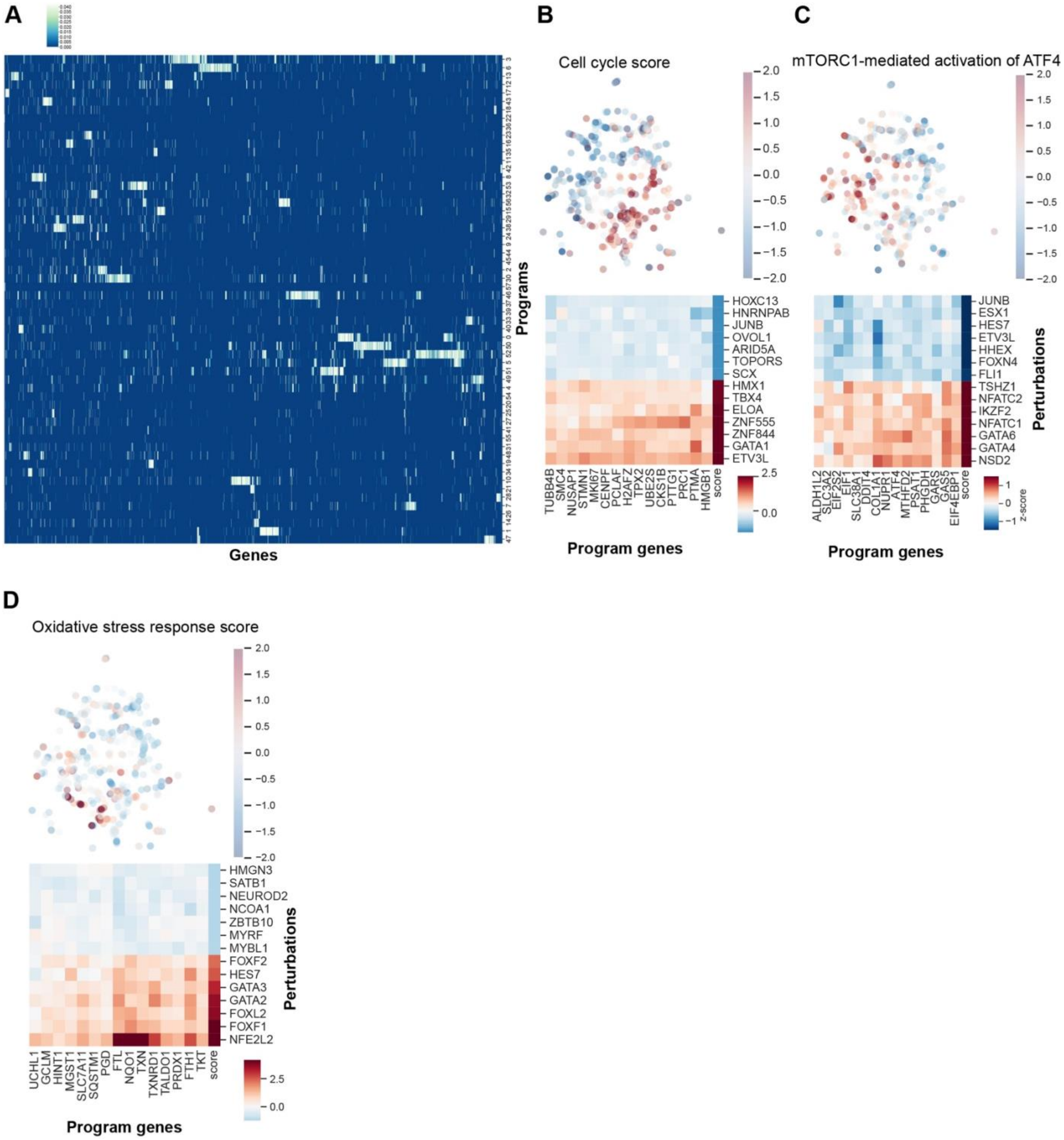


**Figure S9. Sparse gene expression programs, related to Figure 4**

A. Extraction of positive, sparse gene expression programs. We used a modified sparse PCA approach that enforces positivity of coefficients so that genes in programs rise or fall together. To identify the number of programs to extract, we fit the model over bootstrapped resamples of the dataset, used HDBSCAN to cluster the extracted programs, and then reran the algorithm on the dataset using the number of HDBSCAN clusters as the dimensionality (Methods). Figure shows the structure of sparse gene expression programs, which can overlap and vary in size.
B. Additional example of a broadly acting gene expression program in Hs27 cells: Cell cycle. Top: program score overlayed on perturbation embedding from Figure 4A. Bottom: The top activators (red) and repressors (blue) as in Figure 4 D,E.
C. Additional example of a broadly acting gene expression program in Hs27 cells: mTORC1-mediated activation of ATF4. Top: program score overlayed on perturbation embedding from Figure 4A. Bottom: The top activators (red) and repressors (blue) as in Figure 4 D,E.

D. Another example of a specific gene expression program was an oxidative stress response program governed primarily by *NFE2L2*. Top: program score overlayed on perturbation embedding from Figure 4A. Bottom: The top activators (red) and repressors (blue) as in Figure 4 D,E.

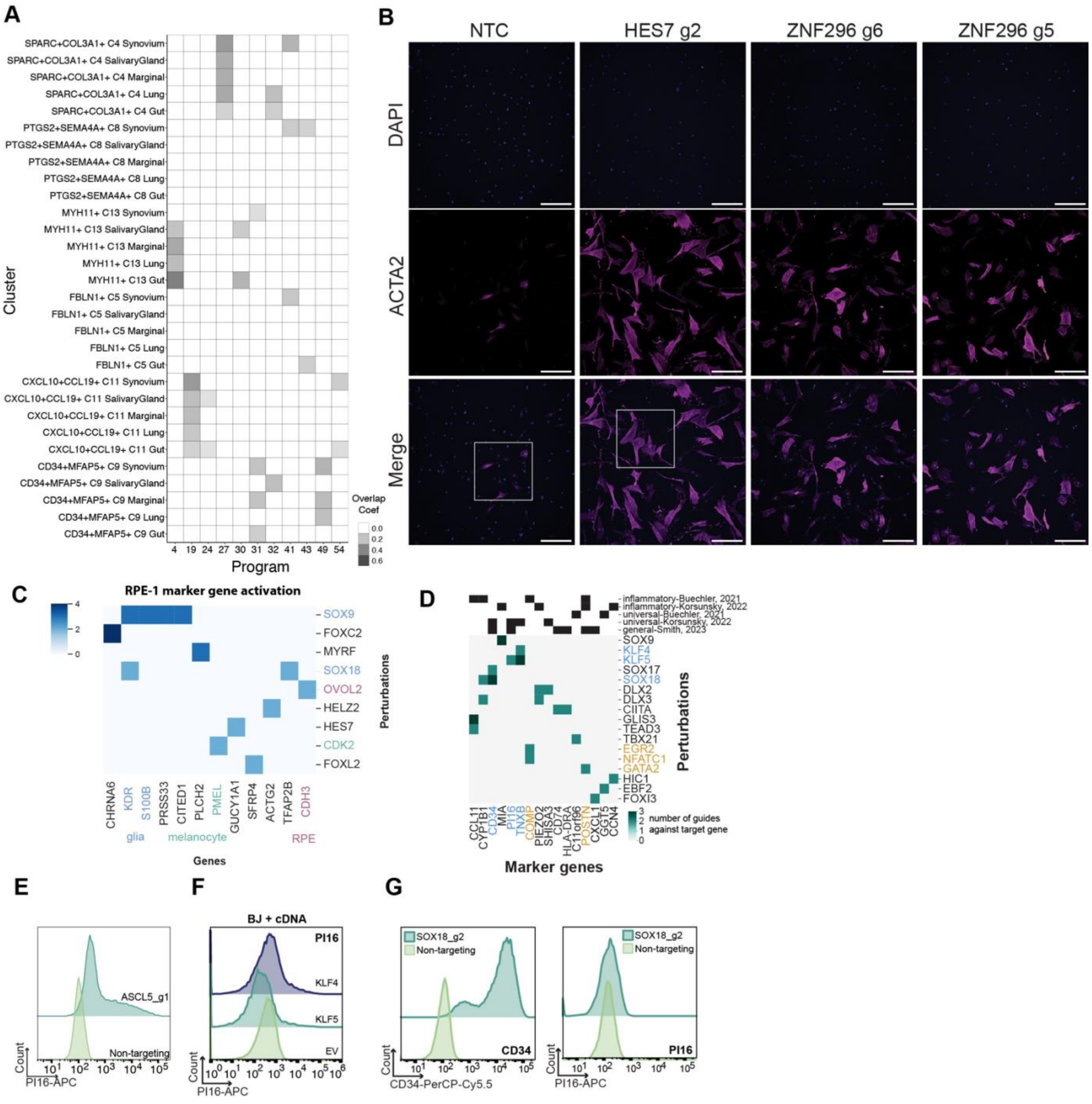


**Figure S10. Studies of newly induced genes, related to Figure 5**

A. Enrichment analysis of gene programs corresponding to *in vivo* cross-tissue fibroblast states from ref. 54. Programs with significant enrichment within each cluster/tissue are colored by their program overlap coefficient.
B. Morphological changes and ACTA2 production by *HES7* and *ZNF296* perturbation. Cells were stained for ACTA2 by immunofluorescence and imaged. Outlines show subsets used in Figure 4D.
C. Heatmap of literature-derived retinal, and related cell type marker genes, that were induced by selected transcription factor perturbations in RPE-1. Marker genes are colored according to the cell type they correspond with (blue: glia, green: melanocyte, red: retinal pigmented epithelial), with perturbations inducing functionally related markers sharing the same color.
D. Heatmap of literature-derived fibroblast marker genes that were induced by selected transcription factor perturbations in Hs27, with sources indicated at top. Marker genes are colored according to the *in vivo* state they correspond with, with perturbations inducing functionally related markers sharing the same color.
E. Flow cytometry validation of PI16 induction by ASCL5_g1 in Hs27 fibroblast cells.
F. Flow cytometry validation of PI16 induction by *KLF4* and *KLF5* cDNA in BJ fibroblast cells.

G. Flow cytometry validation of CD34 induction by SOX18_g2 (left) and corresponding lack of PI16 induction (right) in Hs27 fibroblast cells.
